# Limits to flow detection in phase contrast MRI

**DOI:** 10.1101/2020.03.20.000638

**Authors:** Nathan H. Williamson, Michal E. Komlosh, Dan Benjamini, Peter J. Basser

## Abstract

Pulsed gradient spin echo (PGSE) complex signal behavior becomes dominated by attenuation rather than oscillation when displacements due to flow are similar or less than diffusive displacements. In this “slow-flow” regime, the optimal displacement encoding parameter *q* for phase contrast velocimetry depends on the diffusive length scale 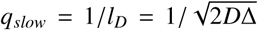 rather than the velocity encoding parameter *v*_*enc*_ = π/(*q*Δ). The minimum detectable mean velocity using the difference between the phase at +*q*_*slow*_ and −*q*_*slow*_ is 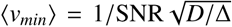. These theories are then validated and applied to MRI by performing PGSE echo planar imaging experiments on water flowing through a column with a bulk region and a beadpack region at controlled flow rates. Velocities as slow as 6 µm/s are detected with velocimetry. Theories, MRI experimental protocols, and validation on a controlled phantom help to bridge the gap between porous media NMR and pre-clinical phase contrast and diffusion MRI.

## 1. Introduction

A single pair of magnetic gradient pulses provides sensitivity to flow through both the signal phase shift and signal attenuation [1]. The phase shift measures the mean velocity of spins within the excited region. The signal attenuation measures the enhanced mixing and spreading in space due to incoherent motion. The lowest detectable velocity is limited by what coherence or added attenuation can be observed in the presence of motion due to self-diffusion.

Slow flows through the brain have recently been identified as part of a mechanism of waste clearance termed glymphatic transport [2]. There is a strong interest to measure slow flows associated with advective transport in the glymphatic system because many neurodegenerative diseases stem from the accumulation of waste products [3, 4]. Researchers have attempted to measure these flows with pulsed gradient spin echo (PGSE) methods [5], however it is still not clear whether these can be detected via MR. Knowledge of the lower limit of velocity resolution, as well as understanding how to best measure slow velocities with PGSE MRI may open the door for imaging glymphatic clearance and other transport processes.

Packer presented the lowest detectable velocity from signal phase in 1969 [6]. We derive a similar equation and provide the optimal sampling strategies for phase contrast velocity measurement in the slow-flow regime. These theories are experimentally validated for MRI applications.

The outline of the paper is as follows. First, a general theory of PGSE NMR is introduced. Then background, literature review, and theory for slow flow measurement with phase contrast velocimetry are provided. Materials and methods are presented to describe the beadpack flow phantom PGSE MRI experiments used for validating the theory and determining the slow flow limits. Methods were tailored towards pre-clinical experiments with considerations towards scan time including the use of PGSE echo planar imaging (EPI) and investigation of the most basic routine for measuring velocity. Then experimental results are provided for phase contrast velocimetry. Significant attention is paid to the phase contrast images. Statistical analysis of regions of interest provide validation of the theory. The Discussion provides suggestions for slow-flow measurements with EPI and *in vivo*, as well as realistic slow flow limits. Then the Discussion focuses on the current understanding of velocities in the glymphatic system, allowing for an assessment of aspects of glymphatic flow which might be measurable with phase contrast MRI.

## 2. Theory

### 2.1. Pulsed gradient spin echo

One beautiful aspect of magnetic resonance is the multitude of information which can be encoded in its complex signal. Pulsed gradient spin echo (PGSE) NMR exemplifies this in the solution to the Bloch-Torrey equations for the normalized complex signal *E*(*q*) arising from a fluid undergoing self-diffusion and coherent flow [1, 7, 8, 9, 10],

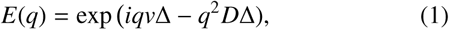

in which *q* sensitizes the signal to displacements during an observation time Δ. With rectangular gradient pulse shapes of amplitude *g* and duration *δ*, in the case that *δ* << Δ, *q* = *γgδ* where *γ* is the gyromagnetic ratio of the nuclei. Equation (1) shows that the phase shift between the real and imaginary channels and the attenuation of the signal intensity provide a means to measure both mean coherent velocity, *v*, and diffusion, *D*, using a single experiment. More generally, the signal describes the sum of the phase shifts associated with molecular displacements, *Z*, during Δ,

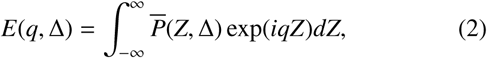

where 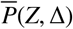 is the average propagator [11]. In the case of Brownian diffusion and coherent flow, the propagator is a Gaussian distribution with mean *v*Δ and standard deviation 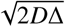,

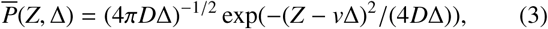

and Eq. (2) becomes equivalent to Eq. (1). Advective (coherent) and dispersive (random) components of mass transport can both be measured with PGSE.

The dispersive component is described by a dispersion coefficient. Flow may not be coherent, i.e. the velocity may be zero when averaged over the voxel. In such situation, the dispersion coefficient may be used as a proxy for the mean velocity ⟨*v*⟩ due to their connection through the Péclet number [12]. The dimensionless Péclet number is the ratio between advective and diffusive mass transport, *Pé* = ⟨*v*⟩ *l*/*D*_0_ where *l* is the characteristic length scale of the flow. Consistent with transport theory, PGSE measurements performed at *Pé* << 1 are sensitive only to diffusion. For *Pé* >> 1, the dispersion coefficient scales with *Pé*. The transition from Brownian motion to flow-enhanced dispersion which occurs near *Pé* = 1 was studied in the longitudinal direction by Ding and Candela [13] and by Kandhai *et al.* [14] and transverse to flow by Scheven [15, 16].

### 2.2. Introduction to phase contrast velocimetry

Coherent velocity is related to the phase, *θ*, the angle between the real and imaginary channels [17, 18], through

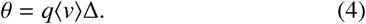

Using the difference between the phase from multiple *q*-points subtracts phase offsets caused by factors other than coherent displacement [19, 20, 21, 22, 23, 24]. This method is called phase contrast velocimetry [25]. The most basic routine for measuring velocity in 1-D is a two-point method [22, 23]:

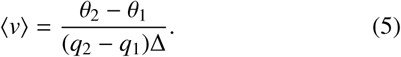

The 3-D velocity vector can be measured using a minimum of 4 *q*-points [25], and Pelc *et al.* have developed and tested such methods [19].

When velocity displacements *v*Δ are similar to diffusive displacements 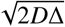, the signal may attenuate with increasing *q* before showing a measurable phase shift. From this standpoint, a simple estimate of the minimum detectable mean velocity is 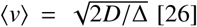 [26]. For reasons which become apparent below, the separation between the slow- and fast-flow regimes is defined as

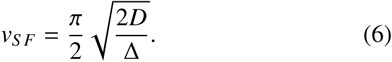

Particularly slow flows have been imaged in plants, where water velocities between 10 and 100 *µ*m/s have been mapped in wheat grain [27], in plant stems [28, 29, 30, 31, 32], in individual giant algae cells [33], in living trees [34, 35], as well as in model soil with the goal of measuring water uptake by roots [36]. Packer developed sequences based on the CPMG which were capable of detecting coherent velocities on the order of 10 *µ*m/s from the buildup of coherent phase shifts [6]. Velocimetry of industrially relevant systems involving slow flows include petrophysics [37, 38], microfluidics [39, 40, 41, 42], chromatography [43, 44], filtration [45, 46] and membrane biofouling [47], and catalytic reactors [48].

In many systems preferential flow directions are known such that displacement encoding gradients and image resolution can be chosen accordingly. Measurement time is not usually limited, and full propagator measurements can be performed to separate mobile and immobile components as well as to obtain mean velocities. Biological systems, in particular mammalian ones, do not provide these luxuries. Likely for these reasons, as well as other confounds, velocimetry in animals has been used mostly to measure velocities in the fast-flow regime. Soon after the first implementation of NMR velocity imaging in humans by Moran in 1982 [21], a number of clinical applications were found [49], including measuring cerebral [25, 50] and aortic [23, 24, 51, 52, 53] blood flow as well as cerebrospinal fluid (CSF) pulsation [54, 55, 56]. One exception was a study of slow velocity fluid flow in tumors by Walker-Samuel *et al.* [57]. They noted that the velocity encoding parameter *v*_*enc*_ (discussed below) needed to be larger than predicted to avoid “crushing” the signal. A better understanding of phase contrast velocimetry measurement in the slow-flow regime could lead to new clinical applications.

Displacements due to coherent flow scale with Δ and eventually become visible with respect to displacements due to random motions, which scale with 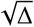. However, the maximum Δ is limited by *T*_2_ (for a spin echo) or *T*_1_ (for a stimulated echo) and is typically somewhere in the window of 10 milliseconds to 1 second for protons in non-viscous fluids. Therefore, in a simple sense, the minimum detectable velocity is limited by the relaxation time and the diffusion coefficient of the nuclei. For a gas, e.g., propane with *D* ∼ 10^−6^ m^2^/s [58], ⟨*v*_*min*_⟩ is on the order of 1 mm/s. Macromolecules and large particles can act as tracers and be sensitive to coherent displacements that are just larger than the particles themselves [59], e.g., for colloids with *D* ∼ 10^−13^ m^2^/s [60] ⟨*v*_*min*_⟩ ∼ 100 nm/s.

For free water, at room temperature with *D* = 2 × 10^−9^ m^2^/s and Δ = 1 s, the simple estimate 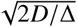 predicts ⟨*v*_*min*_⟩ ≈ 60 *µ*m/s. However, Magdoom *et al.* recently measured velocities of water as slow as 1 *µ*m/s [61]. In practice, the simple estimate significantly over-estimates the true experimental ⟨*v*_*min*_⟩. This is because ⟨*v*_*min*_⟩ is also dependent on the signal-to-noise ratio (SNR), as a few researchers have shown [6, 59, 61].

### 2.3. Limits to flow detection with phase contrast velocimetry

Detection limits and the optimal *q* for phase contrast velocimetry in the slow-flow regime are now derived. The limit to flow detection with Eq. (5) is equivalent to the limit associated with taking the mean of the propagator in the case that the propagator is symmetric about the mean displacement [38, 62]. Well-known detection limits for phase contrast velocimetry in the fast-flow regime are also provided [19].

When the limitation is viewed as the ability to resolve the signal phase shift above the noise floor, connection to the signal-to-noise ratio (SNR = *E*(*q*) |/σ_*noise*_) becomes apparent. Fig. 1 shows complex signal simulated from Eq. (1) with the addition of zero-mean Gaussian noise such that SNR = 100. *q* is non-dimensionalized by the diffusive length scale 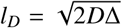. *v* is non-dimensionalized by *v*_*S F*_. At *v*/*v*_*S F*_ = 0.01 the imaginary signal lobes are slightly greater than the standard deviation of the noise, indicating that this velocity can be resolved with SNR = 100. With increasing velocity, the heights and depths of the imaginary lobes increase such that velocity can be measured in the presence of noise with increasing precision.

**Figure 1:**
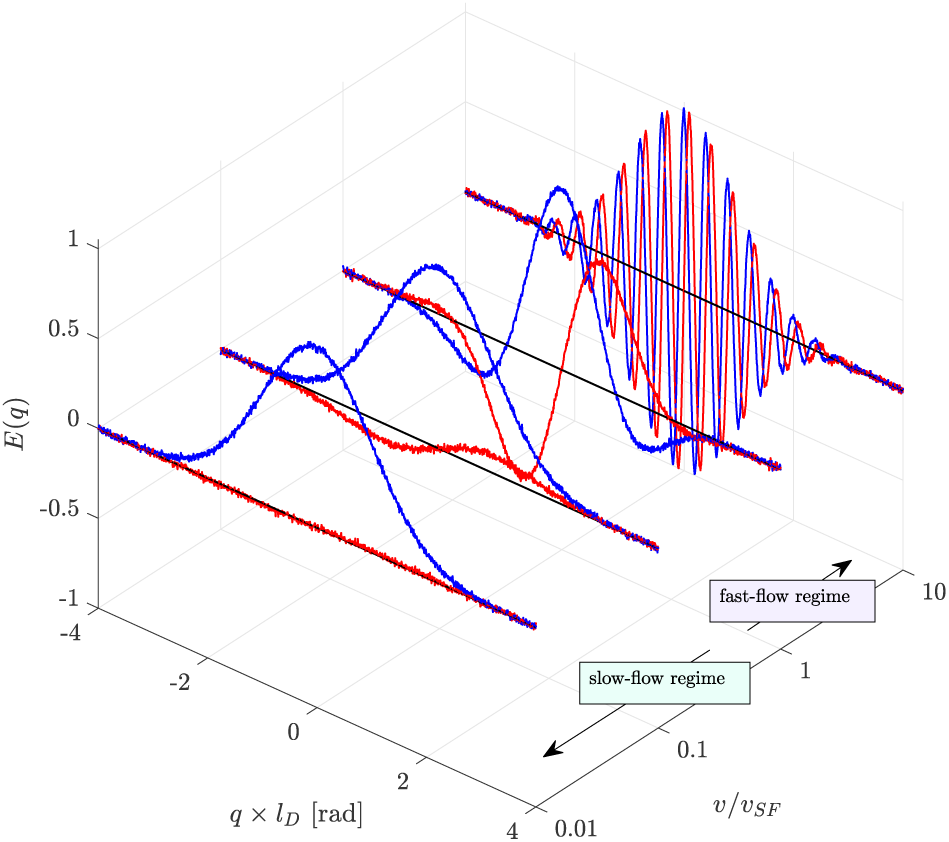
Complex PGSE signal of fluid undergoing flow and diffusion. Real (blue) and imaginary (red) signal simulated from Eq. (1) with SNR = 100 at variable velocities, non-dimensionalized by *l*_*D*_ and *v*_*S F*_.

*v*/*v*_*S F*_ = 1 separates the slow- and fast-flow regimes. In the fast-flow regime, multiple imaginary lobes are resolvable. The imaginary signal lobes peak at *q* = ±π(1 + 2*N*)/(2*v*Δ). N=0 defines the *q* of the first lobe in the fast-flow regime:

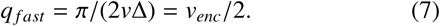

Flow aliasing (wrapping of the phase by intervals of π) occurs when *q* > 2*q*_*f ast*_. This defines the velocity encoding parameter *v*_*enc*_ = π/(*q*Δ) traditionally used in phase contrast MRI [19]. *v*_*enc*_ is chosen to maximize the dynamic range (−*v*_*enc*_ < *v* < +*v*_*enc*_) while maintaining sensitivity to smaller velocities. The standard deviation of the phase from a single *q*-point equals 1/SNR [63]. Pelc *et al.* showed that the standard deviation of velocity measured from two *q*-points is 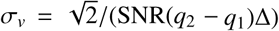 [19]. Therefore, in the fast-flow regime and with *q*_1_ = −*q*_2_ = *q*_*f ast*_, the minimum detectable velocity for a given *q*_*f ast*_ or *v*_*enc*_ is

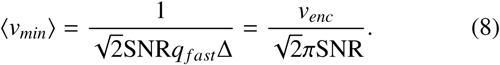

Thus, in the fast-flow regime, measuring slow flows requires smaller *v*_*enc*_ and/or higher SNR [61]. The necessity for smaller *v*_*enc*_ does not hold in the slow-flow regime.

In the slow-flow regime, diffusive attenuation shifts the first imaginary lobe inwards and attenuates the signal before additional lobes are resolved. The imaginary signal is most resolvable above a noise floor at the peak of the first lobe, and thus this corresponding value of *q* provides the greatest sensitivity to flow.

From Euler’s formula, the imaginary component of Eq. (1) is

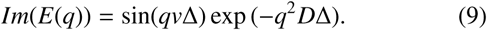

The zero of the first derivative,

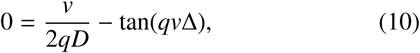

provides the maximum of the first imaginary lobe. This is valid for both the slow- and fast-flow regimes. In the slow-flow regime, tan(*qv*Δ) is approximately *qv*Δ, the first term in a Maclaurin series expansion. Therefore, the imaginary component peaks at approximately

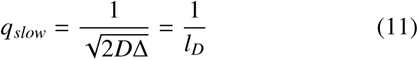

where *l*_*D*_ is the diffusive length scale. The optimum two-point method for 1-D velocimetry of slow flows involves phase contrast measurements at +*q*_*slow*_ and −*q*_*slow*_. No prior knowledge of velocity is needed. At *q*_*slow*_ the signal is attenuated to exp(−1/2) ≈ 0.6 when *δ* << Δ.

The slowest detectable velocity for a given SNR is the velocity at which the imaginary signal is equal to σ_*noise*_. Eq. (11) and (9) can be combined and set equal to 1/SNR to provide

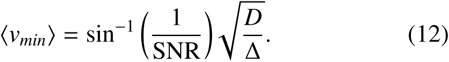

⟨*v*_*min*_⟩ is divided by 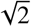 to take into account the doubling of the phase difference but the doubling of the variance when using the two-point method (as in Eq. 8). With SNR >> 1, by series expansion, ⟨*v*_*min*_⟩ simplifies to

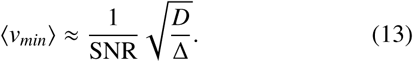

The simple estimate provided by 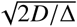 will overestimate ⟨*v*_*min*_⟩ by a factor of 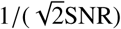.

The numerical solution to Eq. (10) for the *q* value associated with the maximum of the first imaginary lobe is presented in Fig. 2a. The solution converges with *q*_*slow*_ as *v*/*v*_*S F*_ approaches zero and converges with *q*_*f ast*_ when *v*/*v*_*S F*_ >> 1. *q*_*slow*_ and *q*_*f ast*_ cross at *v*/*v*_*S F*_ = 1. Below *v*/*v*_*S F*_ = 1, *q*_*f ast*_ overestimates the *q* necessary for slow-flow measurement and *q*_*slow*_ should be used. Above *v*/*v*_*S F*_ = 1, the situation is reversed and parameters are more appropriately defined using *q*_*f ast*_. Flow aliasing will occur when *q*_*slow*_ is used to measure velocities *v*/*v*_*S F*_ > 2.

**Figure 2:**
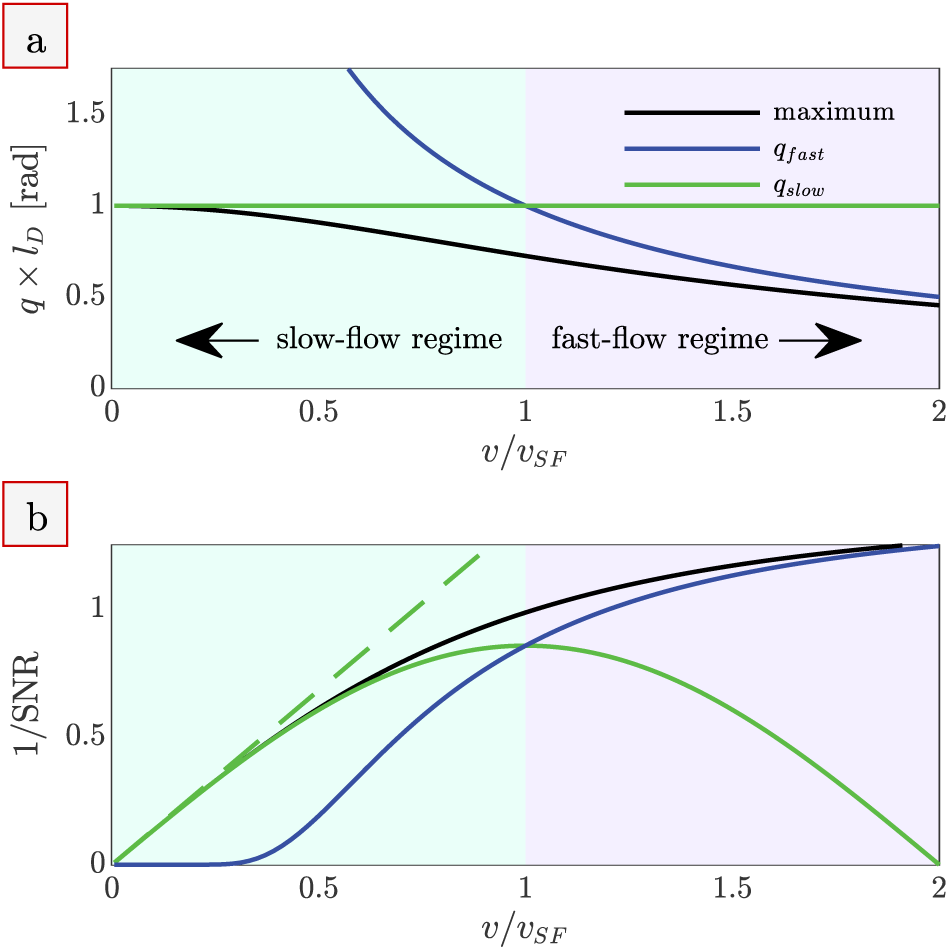
Phase contrast velocimetry in the slow- and fast-flow regimes. (a) The *q* at which the imaginary signal is maximized (numerical solution to Eq. (10), black line), compared to the limiting cases *q*_*slow*_ (Eq. (11), green line) and *q* _*f ast*_ (Eq. (7), blue line). The axes are non-dimensionalized using the diffusive length scale *l*_*d*_ and the velocity separating the slow and fast flow regimes *v*_*S F*_ .n (b) The 1/SNR necessary to detect a given *v*/*v*_*S F*_ when using the *q* from (a) and the two-point method. The dashed green line is the simple expression provided by Eq. (13), valid for SNR >> 1.

Velocity is detectable when the difference in imaginary signal between *E*(*q*_1_) − *E*(*q*_2_) is greater than the standard deviation of the noise. This defines the 1/SNR necessary to detect a given *v* when using the two-point method. The 1/SNR necessary for detection of velocities *v*/*v*_*S F*_ when using *q*_*slow*_, *q*_*f ast*_, and the *q* associated with the maximum of the first imaginary lobe are compared in Fig. 2b. The optimal *q* can be chosen based on *q*_*slow*_ in the slow-flow regime and based on *q*_*f ast*_ or *v*_*enc*_ in the fast-flow regime. However, velocity measurements become imprecise when *q*_*slow*_ or *q*_*f ast*_ are used outside of their regimes. Detection of *v*/*v*_*S F*_ = 0.1 requires SNR > 7 when *q*_*slow*_ is used, but SNR > 1 ×10^19^ when *q*_*f ast*_ (or *v*_*enc*_) is used.

Note that the derivations and Fig. 2 are valid under simplifying assumptions of a shifted Gaussian propagator resulting from a time-independent diffusion coefficient and coherent flow at a single velocity. Additional considerations may be necessary in other flows such as those involving restriction, acceleration, pulsatility, multiphase flow, etc.

### 2.4. Interstitial and superficial velocity

It is necessary to define the average velocity which phase contrast velocimetry measures. The interstitial velocity ⟨*v*⟩ is averaged only over the cross-sectional area taken up by the interstitial fluid; 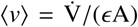 where 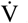 is the flow rate, *ϵ* is the porosity, and A is the entire cross-sectional area. The superficial velocity ⟨*v*_*s*_⟩ is the velocity of the flowing liquid phase averaged over the entire cross-sectional area, ⟨*v*_*s*_⟩ = 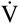 /A. The two averages are related through ⟨*v*_*s*_⟩ = *ϵ* ⟨*v*⟩.

PGSE is sensitive to all proton-bearing fluid in the system. In the case of a beadpack, fluid is only in the interstitial space and PGSE measures the interstitial velocity. In the case of tissue, extracellular (interstitial) water and intracellular water together fill the entire cross-section, and PGSE measures the superficial velocity. The SNR between the two cases of a porous beadpack and a tissue are related through *ϵ*. The superficial velocity of the tissue is less than the interstitial velocity of the beadpack by a factor of *ϵ*, but the SNR is greater by a factor of 1/*ϵ*. The detection limits for these two conditions are theoretically equal.

### 2.5. Phase errors

Experimental phase errors may limit the detection of slow velocities. Many systematic phase errors will cancel out upon taking the phase difference between +*q* and −*q* images (or with whatever encoding scheme is used). For instance, magnetic field inhomogeneity will cause phase errors which are equal for both +*q* and −*q* and hence will cancel out. Phase errors which do not cancel out will lead to an additional phase shift which appears as velocity. Gradient hardware imperfections are a common source of such velocity errors.

For example, magnetic eddy currents induced in surrounding conductive material when ramping gradients lead to unwanted gradients persisting after gradient pulses. Eddy currents can depend on gradient preemphasis calibration and gradient pulse shapes. Persistent unwanted gradients can cause phase shifts [64]. Additionally, mechanical vibrations of the gradients can cause motion of the imaging volume [65]. Pulsed gradients result in concomitant fields which cause phase shifts [66].

Slice and crusher gradients which are in the same direction as the velocity encoding gradients can add or subtract from flow sensitization depending on whether the velocity encoding gradients and slice/crusher gradients are pulsed with the same or reversed polarity. With slice and crusher gradients using positive gradient pulses this unintentionally result in partial flow compensation for −*q* but increased flow sensitivity for +*q*.

Phase errors can differ depending on the location within the sample [66]. Certain corrections may be possible depending on the situation. For example, a mask of phase errors for the entire image can be made by interpolating between pixels known to have no flow [66, 67]. Magdoom *et al.* presented and discussed a stimulated echo (SE) sequence as being beneficial for overcoming phase error introduced by gradient hardware imperfections [61]. Phantoms for which there is known to be no flow can be imaged to determine the extent of the phase errors and to optimize imaging parameters to reduce those errors.

## 3. Materials and methods

A beadpack flow phantom was made from polystyrene microspheres with a 10.03 ± 0.09 *µ*m diameter (Duke Standards, 4000 Series Monosized particles, Thermo Fisher Scientific, USA) packed in water in a 5 mm inner diameter Tricon 5/100 Column (GE Healthcare, USA). Packing occurred by combined sedimentation and a steady flow of 100 *µ*l/min through the column. The porosity *ϵ* = 0.4 was calculated by minimizing the difference between measured flow rates through the bulk and beadpack regions. This beadpack was used as a phantom of glymphatic flow through the brain interstitium, with the bead size chosen to approximate the cell size [68]. The column contained a region of bulk water above the beadpack to mimic paravascular flow. Rigid 1/16” PEEK tubing was used to connect a pump to the phantom and the phantom to the drain. The experimental setup is shown in Fig. 3. A syringe pump (Harvard Apparatus, USA) was used for experiments at flow rates 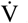 = [0, 0.6, 1, 2, 3, 6, 10, 20, 30, 60, 100, 100, 200, 300] *µ*l/min. The maximum flow rate corresponds to Reynolds number Re ≈ 4 for the bulk region and Re ≈ 0.02 (i.e., creeping flow) for the beadpack region. Flow was turned on or varied at least 10 minutes prior to the start of an experiment.

**Figure 3:**
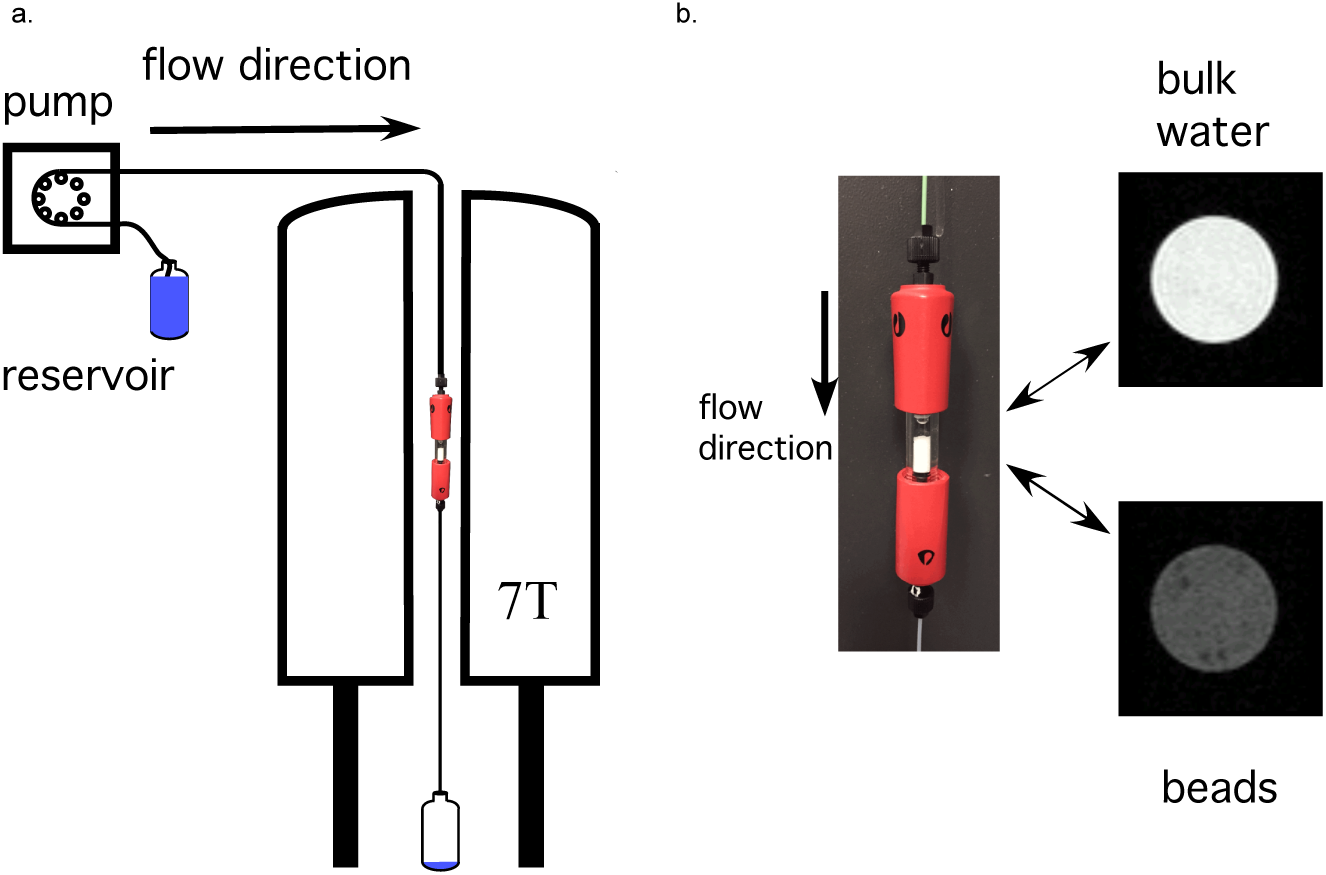
Experimental setup. a) Schematic of flow system with the beadpack flow phantom inside the 7 T vertical bore magnet. b) Picture of beadpack flow phantom and proton density weighted MR images of the bulk water and beadpack region.

MRI experiments were performed with an Avance III spectrometer, a 7T vertical wide-bore magnet, a micro2.5 gradient set and three-axis GREAT 60 amplifiers resulting in a nominal gradient strength of 24.92 mT/m/A (Bruker BioSpin, Germany). The magnet bore temperature was 16.8 °C. The diffusion coefficient in the bulk and beadpack regions of the phantom were measured to be *D*_0_ = 1.89 and 1.2 × 10^−9^ m^2^/s respectively. The setup is shown in Fig. 3. Image analysis was performed in MATLAB R2020a (MathWorks, USA). All experimental data and in-house analysis routines are provided as supplementary material.

A standard diffusion tensor imaging with echo planar imaging (DTI–EPI) sequence was used with Δ/*δ* = 25/2.69 ms, diffusion gradients played out in the *z* direction, and TE/TR=34/2000 ms. 3D images were obtained with double sampling, 8 EPI segments, 1.5 × 1.5 × 2.2 cm field of view and 0.234 × 0.234 × 0.688 mm resolution. No crusher gradients were used. Table 1 shows the z diffusion gradient and *q* values used, including *q* × *l*_*D*_ where *l*_*D*_ = 7.75 *µ*m for the beadpack region. *q* × *l*_*D*_ can be calculated for the bulk region by using *l*_*D*_ = 9.72 *µ*m. *v*_*enc*_ values are also shown. For reference, *q*_1_ and *q*_5_ correspond to *b* = 69 and *b* = 900 mm^2^/s respectively. At each flow rate, an experiment was performed using the positive (+*q*) values and then a second experiment was performed with the same values set negative (−*q*). Each experiment was 42.7 min long.

**Table 1:** z gradients and associated *q* and *v*_*enc*_ values. *q* × *l*_*D*_ uses the diffusive length scale of water in the beadpack.

| Point | $g$ [G/cm] | $q$ [rad/$\mu\text{m}$] | $q \times l_D$ | $v_{\text{enc}}$ [$\mu\text{m}/\text{s}$] |
| --- | --- | --- | --- | --- |
| $q_1$ | 7.46 | 0.0537 | 0.416 | 2,340 |
| $q_2$ | 11.9 | 0.0859 | 0.665 | 1,460 |
| $q_3$ | 17.9 | 0.129 | 0.998 | 975 |
| $q_4$ | 22.4 | 0.161 | 1.25 | 780 |
| $q_5$ | 26.9 | 0.193 | 1.50 | 650 |

For each set of +*q* and −*q* experiments, 3-D magnitude and phase images were reconstructed. A mask was made from the +*q*_1_ magnitude image based on a signal threshold of roughly 12% the maximum voxel signal. The mask was used to omit voxels not within the flow column for magnitude and phase images from the same experiment set.

Phase difference images were calculated from the difference between +*q* and −*q* phase images. A simple unwrapping algorithm was developed where +π was added to voxels with a phase less than 65% of the median of all bordering voxels. The unwrapping algorithm was looped multiple times in order to correct clusters of wrapped voxels as well as voxels which were wrapped by −2π. Then the same was done to unwrap by −π, but only one loop was needed.

z velocity images were calculated from the phase difference images using Eq. (5). Beadpack and bulk regions of interest (ROI) were defined. Both ROIs encompass voxels within three axial slices (shown by dotted lines in Fig. 4). SNR was calculated as the mean signal magnitude from these ROIs divided by the standard deviation of signal magnitude from a region outside of the flow column. The SNR for [*q*_1_, *q*_2_, *q*_3_, *q*_4_, *q*_5_] was [70, 70, 70, 60, 50] for the beadpack and [200, 200, 170, 130, 80] for the bulk, similar for both +*q* and –*q* and for all flow rates.

**Figure 4:**
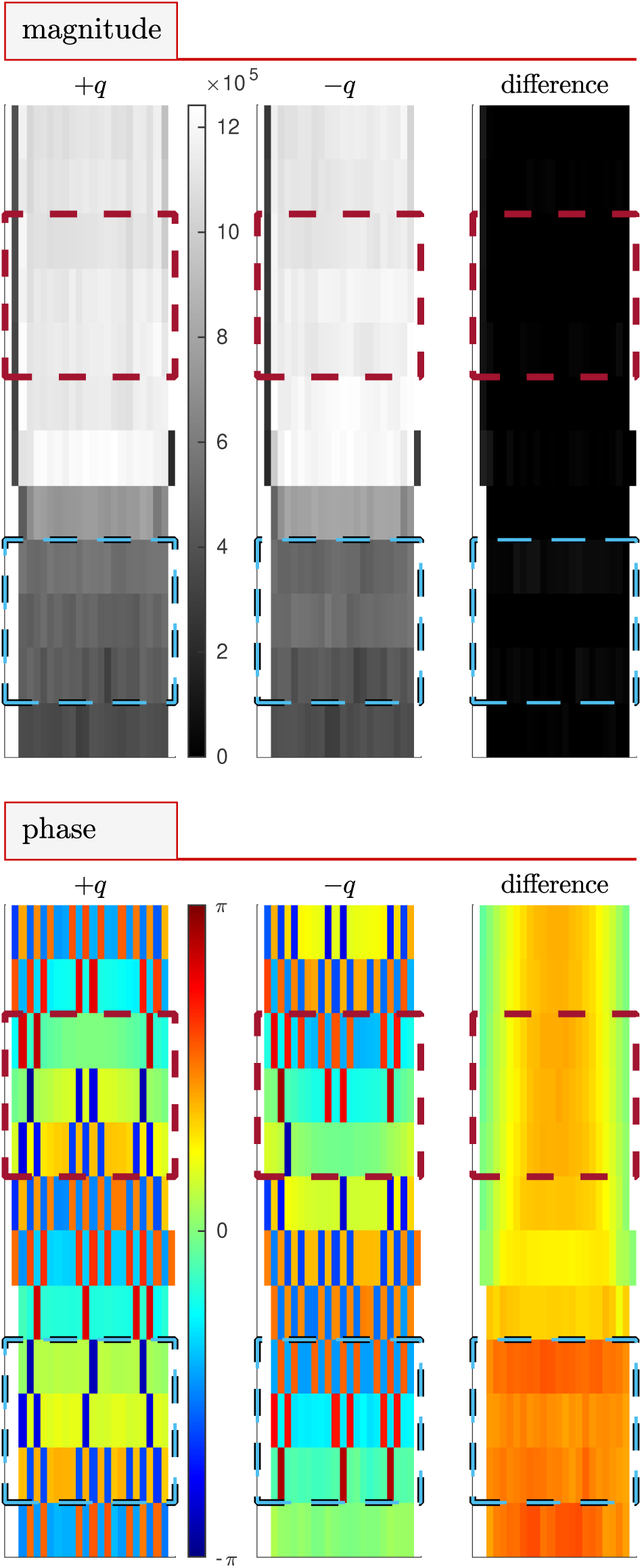
Magnitude and phase images. Sagittal images of magnitude and phase at +*q*_4_, −*q*_4_ and the difference between +*q*_4_ and −*q*_4_ for flow rate 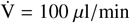 = 100 *µ*l/min. Signal magnitude clearly differentiates Beadpack (dark) and bulk (light) regions. Blue and red dotted lines show the slices included in beadpack and bulk ROIs respectively. Water flow is directed downwards relative to the image.

## 4. Experimental results

### 4.1. Phase contrast MRI in the slow-flow regime

This Section presents the behavior of the phase and velocity for images acquired at variable flow rates in the slow-flow regime. Figures are presented using ±*q*_4_ because this is found minimize the velocity offset (discussed below). MRI using additional *q* and at additional flow rates are presented in the supplementary figures Fig. S1–S11.

Sagittal magnitude and phase images are shown in Fig. 4 for a representative flow rate of 100 *µ*l/min and ±*q*_4_. In magnitude images, the beadpack region is less intense than the bulk region due to the decreased volume fraction of water. Signal magnitude is similar at +*q*_4_ and −*q*_4_ as indicated by the near-zero intensity of the difference image. Phase images appear pixelated an uninformative. However, the phase difference image appears well-behaved. Additional saggital and axial images of magnitude and phase for measurements with and without flow show similar behavior (Figs. S1-S5). Phase difference images acquired at 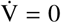 = 0 (Fig. S6) show phase offsets ranging between π/16 < *θ* < +π/16 which depend on *q* and location within the image. Phase offsets lead to velocity offsets when the actual velocity is zero. It is therefore important that the phase offset be near zero.

Sagittal and axial velocity images and velocity profiles for representative flow rates in the low-flow regime and *q*_4_ are presented in Fig. 5. The magnitude images in Fig. 4 can be used as reference for identifying the beadpack and bulk regions. Nonzero velocity offsets in the 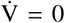 images are due to phase offsets. The velocity offset is particularly apparent when comparing the experimental and theoretical velocity profiles. The offset is constant with increasing flow rate. Therefore, the velocity offset can be corrected in images acquired with flow by subtracting the phase images acquired at 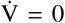, as shown in Fig. S8). Although this is viable for a flow phantom it is not possible for most systems. At faster flow rates the velocity offset is less significant and the measured and theoretical velocity profiles appear similar (shown in Fig. S9).

**Figure 5:**
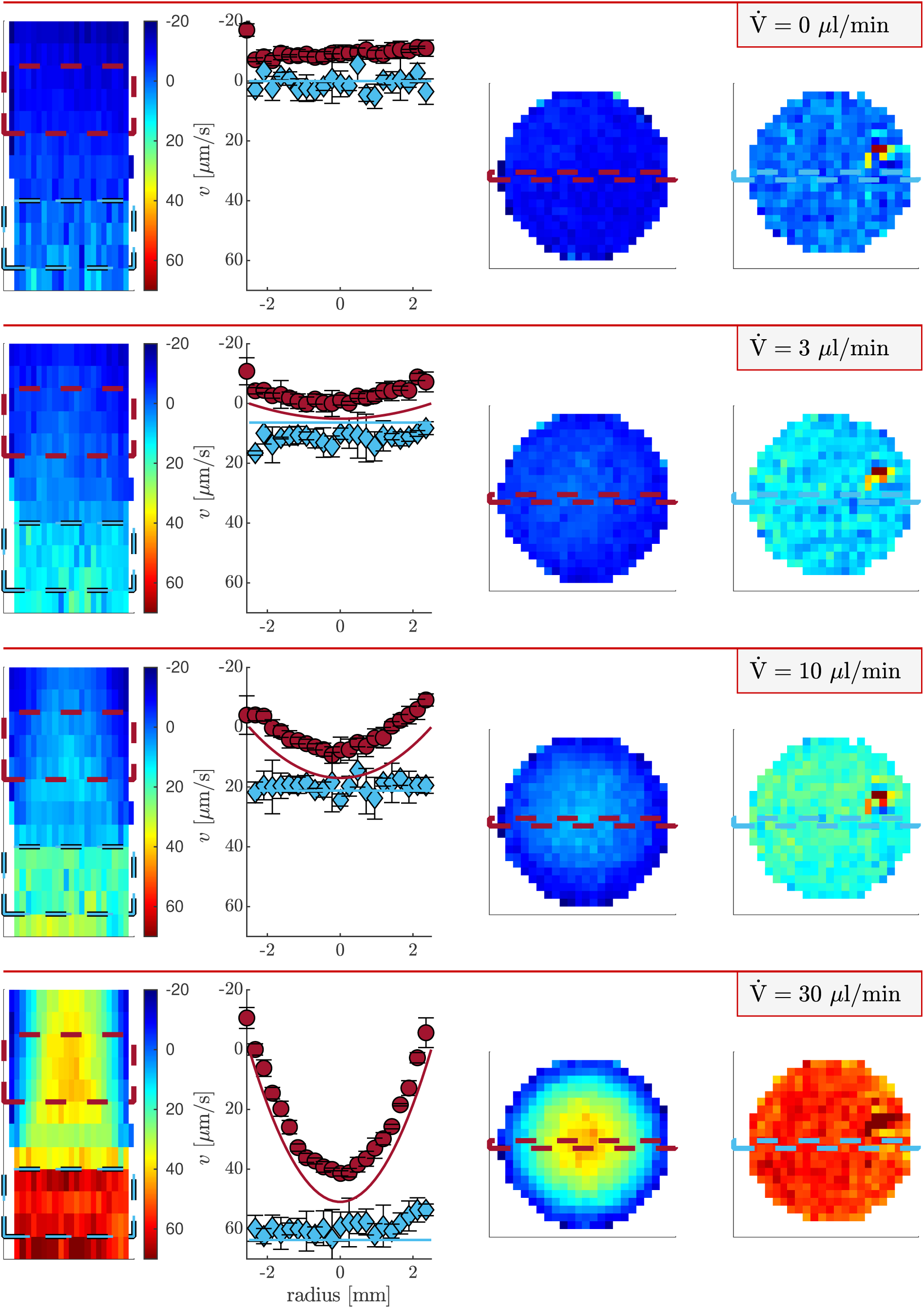
Velocity images and profiles of the beadpack flow phantom. Heat maps of z velocity using *q*_4_ at variable flow rates. Columns 1, 3, and 4 show sagittal, axial bulk, and axial beadpack images respectively. Column 2 shows velocity profiles for the bulk region (red circles) and beadpack region (blue diamonds). The locations from which the profiles are taken are shown by the dashed lines in the axial images. Symbols and error bars are means and standard deviations of three rows of pixels shown by the dashed lines in the saggital image. Solid lines are theoretical velocity profiles for Poiseuille flow and plug flow using the known flow rate, porosity, and tube diameter. 8

A hot spot is seen in the upper right region of axial beadpack images at zero and all flow rates. The hot spot is also seen in the magnitude images (Figs. S3 and S5). The presence of the hot spot at zero flow indicates that it is an imaging artifact.

With increasing flow rate, a parabolic velocity profile develops in the bulk region, as expected for Poiseuille pipe flow. The profile would be expected to deviate and be more blunted than a parabolic profile with increasing proximity to the beadpack interface. This deviation can be seen in the slice just above the beadpack in the sagittal image at 30 *µ*l/min and becomes more apparent at faster flow rates (shown in Fig. S9). This is related to a hydrodynamic entrance length effect which scales with Reynolds number and pipe diameter. The hydrodynamic entrance length of the beadpack is on the order of the bead size and is thus not present in slices below the bulk-beadpack interface. Velocity in the beadpack region is greater than in the bulk region due to the decreased cross-sectional area taken up by the interstitial fluid. The beadpack velocity profiles are relatively constant (plug flow) due to the random packing of the small beads leading to homogeneous coherent displacements when averaged over the length scale of the voxel. At faster flows, a ring of slower velocity is seen at the edge of the tube (Fig. S9).

Although shifted to lower values than theoretically predicted, a parabolic velocity profile is observed for the bulk region at 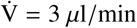. The difference between the velocity at the center and the velocity at the edge is 7 *µ*m/s and is statistically significant. The standard deviation of velocity from groups of three voxels adjacent in the axial direction is 2 *µ*m/s.

### 4.2. Experimental validation of the limits to flow detection

ROI statistics provide a more quantitative perspective on the lowest experimentally detectable velocity. Fig. 6 shows the mean and standard deviations of velocities from all voxels within the beadpack ROI for *q*_3_ and compares them to the theoretically predicted (actual) velocities, assuming plug flow. (See Fig. S10 for other *q* values.) The measured velocities are overall consistent with the actual velocities. In the fast-flow regime, flow aliasing wraps velocity measurements to lower values.

**Figure 6:**
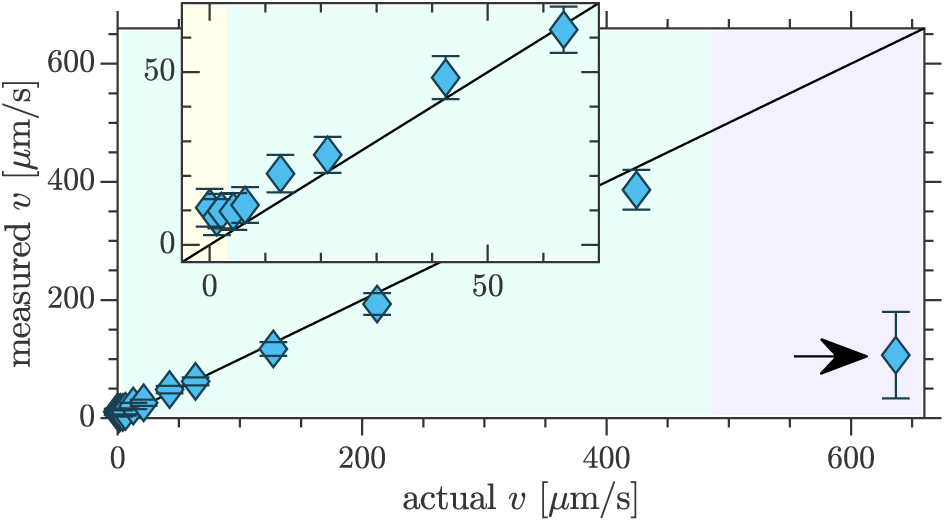
Experimental limits to flow detection with phase contrast velocimetry. Comparison of experimentally measured and actual velocities for flow through the beadpack using *q*_3_. Symbols and errorbars show means and standard deviations from the beadpack ROI defined in Fig. 4. Inset zooms in on the transition from unmeasurable to measurable velocities. Shaded regions show the transitions from theoretically unmeasurable velocities (pale yellow) to the slow-flow regime (pale teal) to the fast-flow regime (pale lilac). Aliasing was observed in the fast-flow regime (marked with arrows).

Statistical significance can be defined as when the difference between the means of the velocity measured at a given flow rate and the velocity measured at zero flow exceed the standard deviation. This first occurs when the actual velocity is 5.8 *µ*m/s at which the measured velocity in the beadpack is 11.8 ± 5.4 *µ*m/s. The theoretical minimum detectable velocity for these conditions is 3.1 *µ*m/s. It is anticipated that there would be additional sources of experimental errors which are not accounted for in the theoretical minimum velocity. Therefore Eq. (13) stands as a close estimate of the lowest detectable velocity with two-point phase contrast velocimetry.

Standard deviations of velocity measurements in the beadpack ROI at zero flow for *q*_1_–*q*_5_ are [6.8, 5.5, 5.5, 5.7, 8.0] *µ*m/s. Theory indicated that in the low-flow regime the standard deviation should be minimized at *q* × *l*_*D*_ = 1. Experimentally, *q*_3_ (*q* × *l*_*D*_ = 0.998) provided the minimum standard deviation, although *q*_2_ (*q* × *l*_*D*_ = 0.665) and *q*_3_ (*q* × *l*_*D*_ = 1.25) were only slightly increased. The approximate similarity between the theoretical and experimental optimum *q* validates Eq. (11).

The *q* should also be chosen to minimize the velocity offset. The velocity offsets for *q*_1_–*q*_5_ are [19, 17, 11, 1.9, 5.0] *µ*m/s, calculated from the mean of velocities in the beadpack ROI at zero flow. *q*_4_ is optimal by a large margin. This indicates that although Eq. (11) provides a rough estimate of the optimum *q* value, it may be necessary to test nearby values to find both the minimum standard deviation and the minimum velocity offset.

The flow rates averaged over each axial slice should equal the pump flow rate. Comparisons of experimentally measured and actual flow rates for the beadpack and bulk ROIs using *q*_3_ are presented in Fig. 7. (Fig. S11 shows similar plots for the other *q* values.) Measured flow rates in the beadpack and bulk ROIs agree well with one another and with the actual flow rate. Deviation at fast flows is due to flow aliasing. Again, the offset at zero flow is apparent and varies with *q* and location within the sample.

**Figure 7:**
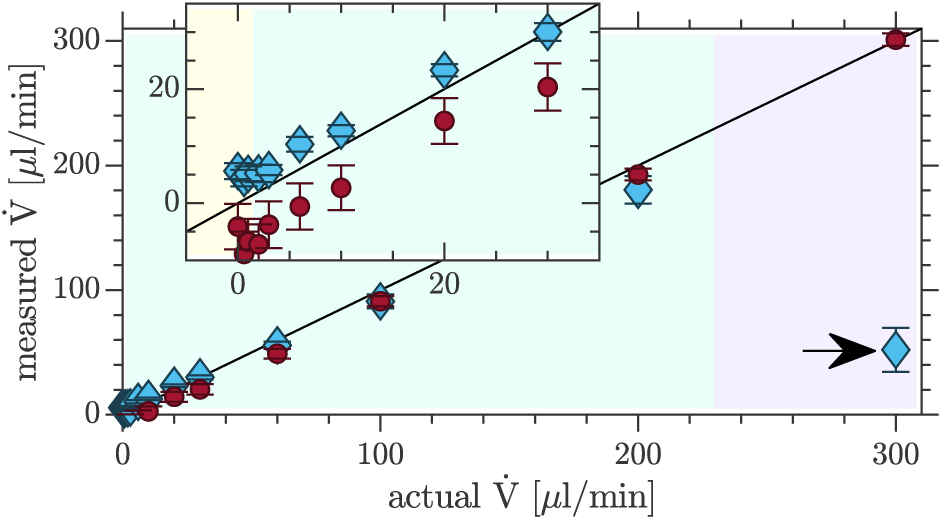
Validation of measured flow rates. Comparison of the actual flow rates to experimentally measured flow rates in the beadpack (blue diamonds) and bulk (red circles) using *q*_3_. Means and standard deviations are from the flow rates measured through three slices for both the beadpack and bulk ROIs, shown in Fig. 4. Shaded regions showing the transitions from theoretically unmeasurable flow rates (pale yellow) to the slow-flow regime (pale teal) to the fast-flow regime (pale lilac) are valid for the beadpack but are slightly different for the bulk (due to different SNR and *D*). Aliasing was observed in the fast-flow regime (marked with arrows).

## 5. Discussion

### 5.1. Phase contrast velocimetry

An offset in phase difference images was found to be dependent on Δ, crusher gradients and slice gradients. Best results were obtained with relatively small Δ, no crusher gradient, and with 3D EPI which minimizes slice gradients. The velocity offset is due to hardware imperfections associated with the slice and crusher gradients, as discussed in Section 2.3. Although theoretically, longer Δ should lead to greater flow sensitivity, experiments indicated that the need to minimize the effective *q* from slice and crusher gradients was more important for the detection of slow flows, and thus Δ =25 ms was used.

### 5.2. Apparent diffusion/dispersion measurement

Dispersion is the spreading of molecules that are initially adjacent in the flow path [12]. The measurement of dispersion with PGSE is well known in the porous media NMR community because it characterizes mass transport without the need for tracers and on length scales which are on the order of the pore size [12, 13, 69, 70, 71, 14, 72, 73, 74, 75, 15]. Diffusion across shear fields, mechanical mixing due to tortuous flow paths around obstacles, and holdup of molecules in stagnant regions all cause dispersion. As discussed in the theory section, the apparent diffusion or dispersion coefficient from signal attenuation can be used as a proxy for velocity.

The slowest velocity which is detectable by diffusion/dispersion measurements is roughly defined by *Pé* = 1 [13, 14]. The value of the slowest velocity for a certain application depends on a characteristic length scale of the flow *l* which is ambiguous for heterogeneous media. Enhanced dispersion is seen when 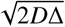 > *l* [70, 73]. Therefore, at best, the minimum detectable velocity with diffusion/dispersion measurement is greater than 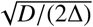. It is seen that the factors limiting the detection of flow with diffusion/dispersion measurement are similar to those affecting phase contrast velocimetry. Under the same conditions, the minimum velocity for phase contrast velocimetry is less than that of diffusion/dispersion measurement.

However, diffusion/dispersion measurements may be advantageous over phase contrast velocimetry due to the magnitude signal being prone to less artifacts than the phase, particularly in the presence of sample motion and when combined with fast imaging methods. Additionally, it may not always be possible to reach sufficiently high resolution to observe coherent flow associated with the process of interest.

Some of the first diffusion MRI studies performed on human brains observed dispersion caused by tissue perfusion [76, 77]. In particular, the microcirculation of blood in capillary networks produced intravoxel incoherent motion (IVIM) in addition to the random Brownian motion of water molecules diffusing through the tissue. Diffusion tensor imaging (DTI) methods have also been developed to assess cerebrospinal fluid motion along perivascular spaces [78, 5] (a key component of glymphatic clearance [79]).

### 5.3. Other PGSE methods

There are alternative experimental and analysis methods based on PGSE which can be useful in certain slow-flow applications. The asymmetric component of the propagator has been used to measure slow flows in plants [80]. There are methods of reducing the data requirements of propagator measurements which are applicable to slow flows [81, 82]. A propagator provides more information about flow than *D*_*app*_ and ⟨*v*⟩ in instances where it cannot be simply described by a shifted Gaussian. This commonly occurs in multiphase flow where there is a slower or stagnant phase and a more mobile phase as well as flow through porous materials where length scales of the material heterogeneity are longer than the length scales of fluid displacement.

This experimental study used pulsed gradient spin echo for velocity encoding. Use of a stimulated echo such that the maximum Δ is limited by *T*_1_ rather than *T*_2_ can be a strategy for detecting slow flows with phase contrast velocimetry [83, 28, 84, 61]. Gradient echo flow encoding has also been used for slow-flow measurement although this significantly reduces the maximum Δ [57].

Alternative pulse sequences may be useful for detecting or isolating slowly moving fluid. Flow compensated double PGSE NMR refocuses signal from spins with locally coherent velocities such that attenuation is only caused by stochastic fluctuations [70, 73]. This experiment has been used to isolate biologically relevant IVIM parameters [85], and may also be useful for detecting local coherence associated with slowly flowing fluid. Sequences built off of the CPMG such as the ones proposed by Packer [6] are able to amplify phase shifts of slowly moving fluid, although combining them with imaging may be difficult.

### 5.4. Limits to flow detection in vivo

The theory above provides useful approximations of slow-flow detection limits for *in vivo* phase contrast MRI. With water *D*_0_ = 3 × 10^−9^ m^2^/s at 37°*C* [86], SNR = 100, and Δ = 1 *s*, velocities as slow as ⟨*v*_*min*_⟩ = 1 *µ*m/s may be detectable with phase contrast velocimetry. The velocity must be coherent on the length scale of the voxel.

Here, phase contrast velocimetry experiments were capable of detecting velocities as slow as 6 *µ*m/s. These experiments can be performed in a pre-clinical setting with very little adjustment. However, here the direction of flow was known *a priori*. Only ±*q*_*z*_ was utilized for phase contast velocimetry and therefore only *v*_*z*_ was measured. (Note that propagator measurements in the *x* and *y* directions showed no measurable coherent *v*_*x*_ or *v*_*y*_ flow in the phantom [68].) Standard phase contrast encoding methods can be combined with the theory here to measure the velocity vector [19]. Based on the significant phase errors caused by slice and and crusher gradients it is worthwhile investigating different encoding strategies. In particular a more conservative six-*q*-point method involving +*q* and −*q* for the three Cartesian directions may subtract more phase errors than the faster four-*q*-point method (sometimes referred to as Hadamard encoding) [87].

Minimum detectable velocities can be compared to published values for the velocities associated with *in vivo* mass transport for a preliminary assessment of future applications. Future developments such as methods which enhance SNR, or which select signal of endogenous molecules with smaller diffusion coefficients or spins with longer relaxation times may be able to push these limits even further and provide opportunity for new low-flow applications.

### 5.5. Preliminary assessment of the applicability of phase contrast MRI to imaging glymphatic clearance

The glymphatic system may play a critical role in prevention and recovery from neurodegenerative diseases and injuries. Currently, there is no clinical measure of glymphatic function. Dynamic contrast enhanced (DCE) MRI methods show promise [88], however intrathecal gadolinium contrast agent injection may be considered too invasive and risky in most cases. PGSE MRI may have the potential to image features of glymphatic clearance without requiring exogeneous contrast agents. For a preliminary assessment, minimum detectable velocities observed with PGSE MRI can be compared to literature values for velocities associated with glymphatic clearance (which were recently compiled in a review article [89]). Note that there are very few experiments in which velocities associated with glymphatic clearance are reported (also highlighting the need for developing quantitative non-invasive velocimetry methods). With much unknown about the glymphatic system, the reported velocities are placeholders for future experimental findings.

In the glymphatic hypothesis, fluid moves into the brain along periarterial spaces, through the brain interstitium, and out of the brain along perivenous spaces [2]. Glymphatic clearance has been shown to increase during sleep [90]. DCE MRI measurements on live rat brains indicate diffusion and advection both contribute to interstitial transport of the contrast agent [91]. This and other research [92] suggests that the *Pé* of the contrast agent is in an intermediate regime (near *Pé* = 1), but that the *Pé* of water is less than 1. Finite element models were found to give the best comparison with real-time iontophoresis experiments of tetramethylammonium transport in brains of anesthetized young adult mice when a superficial velocity of ⟨*v*_*s*_⟩ = 10 *µ*m/s was included [92]. An upper bound for superficial velocity was estimated at ⟨*v*_*s*_⟩ < 50 *µ*m/s. During wakefulness, ⟨*v*_*s*_⟩ < 1 *µ*m/s was estimated.

In tissue, PGSE measures a mean velocity which is similar to the superficial velocity ⟨*v*_*s*_⟩. Therefore, detection limits can be compared to the ⟨*v*_*s*_⟩ values quoted above. Phase contrast velocimetry may be able to measure interstitial glymphatic flow. High resolution would be necessary to avoid the influence of partial volume effects from blood and CSF and for directed flow to not be averaged out. In particular, voxel sizes may need to be smaller than the distance between arterioles and venules, which has been measured at roughly 175 *µ*m in mice [93] and 280 *µ*m in primates [94].

Perivascular CSF flow is another point in the proposed mechanism of glymphatic clearance to which PGSE MRI might be sensitive. CSF in the annular spaces between vascular walls and astrocytic endfeet is connected to CSF in the subarachnoid space and the ventricles which bathes the brain [95, 96]. Tracer movement into the brain occurs along periarterial spaces faster than would be predicted by diffusion alone [97]. Pulsation of arterial walls causes motion of CSF [95, 79]. The glymphatic hypothesis is that there is peristaltic flow of CSF in the direction of blood flow and that the flow continues out of the brain on the perivenule side [2, 79]. Particle tracking velocimetry of fluorescent microspheres by two photon spectroscopy measured pulsatile flow of CSF in periarterial spaces of mice with velocities between 17 − 19 *µ*m/s in the direction of blood flow [98, 99]. These velocities are averaged over many cardiac cycles. The velocity profile was parabolic, as expected for laminar flow [98]. Microspheres in periarterial spaces of mice oscillated back and forth Δ*x* = 14 ± 2*µ*m per cycle with approximately 360 bpm heart rate [99]. Instantaneous velocities reached greater than 100 *µ*m/s. These velocities could be detectable with phase contrast velocimetry. Perivascular spaces were seen to be 40 *µ*m wide [98]. Although such high resolution can be difficult to achieve *in vivo*, there are image processing approaches to enhance perivascular space conspicuity and quantification [100].

## 6. Conclusion

Velocity detection with single pulsed gradient spin echo (PGSE) NMR is known to be limited by the smearing of the phase coherence by diffusion and signal loss by dispersive attenuation. Detection limits and the optimal *q* for phase contrast velocimetry in the slow-flow regime are derived. The theory points out that the *v*_*enc*_ parameter is intended for the fast-flow regime and can lead to the signal being attenuated into noise if used in the slow-flow regime. Theories are experimentally validated with PGSE EPI measurements on a beadpack flow phantom using methods available in a pre-clinical setting. The lowest detectable velocities *in vivo* are discussed and are used to preliminarily assess the applicability of PGSE MRI to imaging glymphatic clearance based on a survey of literature values for glymphatic flows. This study is intended to aid in the translation of porous media NMR concepts to applications of slow-flow measurement in pre-clinical phase contrast and diffusion MRI.

## Acknowledgements

NHW was funded by the NIGMS PRAT fellowship Award # FI2GM133445-01. PJB was supported by the IRP of the NICHD, NIH. DB and MK were supported by the Center for Neuroscience and Regenerative Medicine, Henry Jackson Foundation, Bethesda, MD. NHW is grateful for mentorship from Prof. Joseph Seymour and Prof. Sarah Codd.

## Supplementary material

The following files have been made available as supplementary material: pdf document containing supplementary figures (Figs. S1–S11); MATLAB code for phase contrast MR image construction and analysis; MATLAB file containing a structure array of the 3D phase and magnitude image data at all flow rates.

## References

[1] E. Stejskal, Use of spin echoes in a pulsed magnetic-field gradient to study anisotropic, restricted diffusion and flow, The Journal of Chemical Physics 43 (10) (1965) 3597–3603 (1965).

[2] J. J. Iliff, M. Wang, Y. Liao, B. A. Plogg, W. Peng, G. A. Gundersen, H. Benveniste, G. E. Vates, R. Deane, S. A. Goldman, et al., A paravascular pathway facilitates csf flow through the brain parenchyma and the clearance of interstitial solutes, including amyloid β, Science translational medicine 4 (147) (2012) 147ra111–147ra111 (2012).

[3] B. A. Plog, M. Nedergaard, The glymphatic system in central nervous system health and disease: past, present, and future, Annual Review of Pathology: Mechanisms of Disease 13 (2018) 379–394 (2018).

[4] T. Taoka, S. Naganawa, Glymphatic imaging using mri, Journal of Magnetic Resonance Imaging 51 (1) (2020) 11–24 (2020).

[5] I. F. Harrison, B. Siow, A. B. Akilo, P. G. Evans, O. Ismail, Y. Ohene, P. Nahavandi, D. L. Thomas, M. F. Lythgoe, J. A. Wells, Non-invasive imaging of csf-mediated brain clearance pathways via assessment of perivascular fluid movement with diffusion tensor mri, Elife 17 (2018) e34028.(2018).

[6] K. Packer, The study of slow coherent molecular motion by pulsed nuclear magnetic resonance, Molecular Physics 17 (4) (1969) 355–368 (1969).

[7] F. Bloch, Nuclear induction, Physical Review 70 (7-8) (1946) 460 (1946).

[8] H. C. Torrey, Bloch equations with diffusion terms, Physical Review 104 (1956) 563–565 (Nov 1956).

[9] E. Stejskal, J. Tanner, Spin diffusion measurements: Spin echoes in the presence of a time-dependent field gradient, The Journal of Chemical Physics 42 (1962) 288–298 (1962).

[10] P. T. Callaghan, C. Eccles, Y. Xia, Nmr microscopy of dynamic displacements: k-space and q-space imaging, Journal of Physics E: Scientific Instruments 21 (8) (1988) 820 (1988).

[11] J. Kärger, W. Heink, The propagator representation of molecular transport in microporous crystallites, Journal of Magnetic Resonance 51 (1) (1983) 1 – 7 (1983).

[12] J. D. Seymour, P. T. Callaghan, Generalized approach to nmr analysis of flow and dispersion in porous media, AIChE Journal 43 (8) (1997) 2096–2111 (1997).

[13] A. Ding, D. Candela, Probing nonlocal tracer dispersion in flows through random porous media, Physical Review E 54 (1) (1996) 656 (1996).

[14] D. Kandhai, D. Hlushkou, A. G. Hoekstra, P. M. Sloot, H. Van As, U. Tallarek, Influence of stagnant zones on transient and asymptotic dispersion in macroscopically homogeneous porous media, Physical review letters 88 (23) (2002) 234501 (2002).

[15] U. Scheven, Pore-scale mixing and transverse dispersivity of randomly packed monodisperse spheres, Physical review letters 110 (21) (2013) 214504 (2013).

[16] U. Scheven, S. Khirevich, A. Daneyko, U. Tallarek, Longitudinal and transverse dispersion in flow through random packings of spheres: A quantitative comparison of experiments, simulations, and models, Physical Review E 89 (5) (2014) 053023 (2014).

[17] E. Hahn, Spin echoes, Physical Review 80 (1950) 580–594 (1950).

[18] H. Y. Carr, E. M. Purcell, Effects of diffusion on free precession in nuclear magnetic resonance experiments, Physical review 94 (3) (1954) 630 (1954).

[19] N. J. Pelc, M. A. Bernstein, A. Shimakawa, G. H. Glover, Encoding strategies for three-direction phase-contrast mr imaging of flow, Journal of Magnetic Resonance Imaging 1 (4) (1991) 405–413 (1991).

[20] T. Grover, J. Singer, Nmr spin-echo flow measurements, Journal of Applied Physics 42 (3) (1971) 938–940 (1971).

[21] P. R. Moran, A flow velocity zeugmatographic interlace for nmr imaging in humans, Magnetic resonance imaging 1 (4) (1982) 197–203 (1982).

[22] D. A. Feinberg, L. E. Crooks, P. Sheldon, J. H. Iii, J. Watts, M. Arakawa, Magnetic resonance imaging the velocity vector components of fluid flow, Magnetic resonance in medicine 2 (6) (1985) 555–566 (1985).

[23] M. O’donnell, Nmr blood flow imaging using multiecho, phase contrast sequences, Medical physics 12 (1) (1985) 59–64 (1985).

[24] G. Nayler, D. Firmin, D. Longmore, et al., Blood flow imaging by cine magnetic resonance, J Comput Assist Tomogr 10 (5) (1986) 715–722 (1986).

[25] C. Dumoulin, S. Souza, M. Walker, W. Wagle, Three-dimensional phase contrast angiography, Magnetic Resonance in Medicine 9 (1) (1989) 139–149 (1989).

[26] P. T. Callaghan, Principles of nuclear magnetic resonance microscopy, Clarendon Press, Oxford, 1991 (1991).

[27] C. Jenner, Y. Xia, C. Eccles, P. Callaghan, Circulation of water within wheat grain revealed by nuclear magnetic resonance micro-imaging, Nature 336 (6197) (1988) 399 (1988).

[28] Y. Xia, P. T. Callaghan, “one-shot” velocity microscopy: Nmr imaging of motion using a single phase-encoding step, Magnetic resonance in medicine 23 (1) (1992) 138–153 (1992).

[29] Y. Xia, V. Sarafis, E. Campbell, P. Callaghan, Non invasive imaging of water flow in plants by nmr microscopy, Protoplasma 173 (3-4) (1993) 170–176 (1993).

[30] W. Köckenberger, J. Pope, Y. Xia, K. Jeffrey, E. Komor, P. Callaghan, A non-invasive measurement of phloem and xylem water flow in castor bean seedlings by nuclear magnetic resonance microimaging, Planta 201 (1) (1997) 53–63 (1997).

[31] M. Rokitta, U. Zimmermann, A. Haase, Fast nmr flow measurements in plants using flash imaging, Journal of Magnetic Resonance 137 (1) (1999) 29–32 (1999).

[32] A. Peuke, M. Rokitta, U. Zimmermann, L. Schreiber, A. Haase, Simultaneous measurement of water flow velocity and solute transport in xylem and phloem of adult plants of ricinus communis over a daily time course by nuclear magnetic resonance spectrometry, Plant, Cell & Environment 24 (5) (2001) 491–503 (2001).

[33] J.-W. van de Meent, A. J. Sederman, L. F. Gladden, R. E. Goldstein, Measurement of cytoplasmic streaming in single plant cells by magnetic resonance velocimetry, Journal of Fluid Mechanics 642 (2010) 5–14 (2010).

[34] C. W. Windt, P. Blümler, A portable nmr sensor to measure dynamic changes in the amount of water in living stems or fruit and its potential to measure sap flow, Tree Physiology 35 (4) (2015) 366–375 (2015).

[35] A. Nagata, K. Kose, Y. Terada, Development of an outdoor mri system for measuring flow in a living tree, Journal of Magnetic Resonance 265 (2016) 129 – 138 (2016).

[36] N. Spindler, P. Galvosas, A. Pohlmeier, H. Vereecken, Nmr velocimetry with 13-interval stimulated echo multi-slice imaging in natural porous media under low flow rates, Journal of Magnetic Resonance 212 (1) (2011) 216 – 223 (2011).

[37] R. Waggoner, E. Fukushima, Velocity distribution of slow fluid flows in bentheimer sandstone: An nmri and propagator study, Magnetic Resonance Imaging 14 (9) (1996) 1085 – 1091 (1996).

[38] M. N. Shukla, A. Vallatos, V. R. Phoenix, W. M. Holmes, Accurate phase-shift velocimetry in rock, Journal of Magnetic Resonance 267 (2016) 43 – 53 (2016).

[39] B. S. Akpa, S. M. Matthews, A. J. Sederman, K. Yunus, A. C. Fisher, M. L. Johns, L. F. Gladden, Study of miscible and immiscible flows in a microchannel using magnetic resonance imaging, Analytical Chemistry 79 (16) (2007) 6128–6134 (2007).

[40] J. Zhang, B. J. Balcom, Parallel-plate rf resonator imaging of chemical shift resolved capillary flow, Magnetic resonance imaging 28 (6) (2010) 826–833 (2010).

[41] S. Ahola, V.-V. Telkki, S. Stapf, Velocity distributions in a micromixer measured by nmr imaging, Lab on a Chip 12 (10) (2012) 1823–1830 (2012).

[42] M. Wiese, S. Benders, B. Blümich, M. Wessling, 3d mri velocimetry of non-transparent 3d-printed staggered herringbone mixers, Chemical Engineering Journal 343 (2018) 54 – 60 (2018).

[43] U. Tallarek, D. Van Dusschoten, H. Van As, E. Bayer, G. Guiochon, Study of transport phenomena in chromatographic columns by pulsed field gradient nmr, The Journal of Physical Chemistry B 102 (18) (1998) 3486–3497 (1998).

[44] J. Park, S. J. Gibbs, Mapping flow and dispersion in a packed column by mri, AIChE Journal 45 (3) (1999) 655–660 (1999). arXiv:https://onlinelibrary.wiley.com/doi/pdf/10.1002/aic.690450322.

[45] L. Huang, G. Mikolajczyk, E. Küstermann, M. Wilhelm, S. Odenbach, W. Dreher, Adapted mr velocimetry of slow liquid flow in porous media, Journal of Magnetic Resonance 276 (2017) 103 – 112 (2017).

[46] G. Mikolajczyk, L. Huang, M. Wilhelm, W. Dreher, S. Odenbach, Colloid deposition in monolithic porous media – experimental investigations using x-ray computed microtomography and magnetic resonance velocimetry, Chemical Engineering Science 175 (2018) 257 – 266 (2018).

[47] D. G. von der Schulenburg, J. Vrouwenvelder, S. Creber, M. van Loosdrecht, M. Johns, Nuclear magnetic resonance microscopy studies of membrane biofouling, Journal of Membrane Science 323 (1) (2008) 37 – 44 (2008).

[48] L. Gladden, M. Lim, M. Mantle, A. Sederman, E. Stitt, Mri visualisation of two-phase flow in structured supports and trickle-bed reactors, Catalysis Today 79 (2003) 203–210 (2003).

[49] N. J. Pelc, R. J. Herfkens, A. Shimakawa, D. R. Enzmann, et al., Phase contrast cine magnetic resonance imaging, Magnetic resonance quarterly 7 (4) (1991) 229–254 (1991).

[50] M. Marks, N. Pelc, M. Ross, D. Enzmann, Determination of cerebral blood flow with a phase-contrast cine mr imaging technique: evaluation of normal subjects and patients with arteriovenous malformations., Radiology 182 (2) (1992) 467–476 (1992).

[51] M. C. Dulce, G. H. Mostbeck, M. O’Sullivan, M. Cheitlin, G. R. Caputo, C. B. Higgins, Severity of aortic regurgitation: interstudy reproducibility of measurements with velocity-encoded cine mr imaging., Radiology 185 (1) (1992) 235–240 (1992).

[52] J.-P. E. Kvitting, T. Ebbers, L. Wigström, J. Engvall, C. L. Olin, A. F. Bolger, Flow patterns in the aortic root and the aorta studied with time-resolved, 3-dimensional, phase-contrast magnetic resonance imaging: implications for aortic valve–sparing surgery, The Journal of Thoracic and Cardiovascular Surgery 127 (6) (2004) 1602 – 1607 (2004).

[53] A. Frydrychowicz, M. Markl, D. Hirtler, A. Harloff, C. Schlensak, J. Geiger, B. Stiller, R. Arnold, Aortic hemodynamics in patients with and without repair of aortic coarctation: in vivo analysis by 4d flow-sensitive magnetic resonance imaging, Investigative radiology 46 (5) (2011) 317–325 (2011).

[54] D. Enzmann, N. Pelc, Normal flow patterns of intracranial and spinal cerebrospinal fluid defined with phase-contrast cine mr imaging., Radology 178 (2) (1991) 467–474 (1991).

[55] D. R. Enzmann, N. J. Pelc, Cerebrospinal fluid flow measured by phase-contrast cine mr, American journal of neuroradiology 14 (6) (1993) 1301–1307 (1993).

[56] A. A. Linninger, M. Xenos, D. C. Zhu, M. R. Somayaji, S. Kondapalli, R. D. Penn, Cerebrospinal fluid flow in the normal and hydrocephalic human brain, IEEE Transactions on Biomedical Engineering 54 (2) (2007) 291–302 (2007).

[57] S. Walker-Samuel, T. A. Roberts, R. Ramasawmy, J. S. Burrell, S. P. Johnson, B. M. Siow, S. Richardson, M. R. Goncalves, D. Pendse, S. P. Robinson, et al., Investigating low-velocity fluid flow in tumors with convection-mri, Cancer research 78 (7) (2018) 1859–1872 (2018).

[58] S. Codd, S. Altobelli, A pgse study of propane gas flow through model porous bead packs, Journal of Magnetic Resonance 163 (1) (2003) 16 – 22 (2003).

[59] H. Wassenius, P. T. Callaghan, Nanoscale nmr velocimetry by means of slowly diffusing tracer particles, Journal of Magnetic Resonance 169 (2) (2004) 250 – 256 (2004).

[60] E. O. Fridjonsson, J. D. Seymour, S. L. Codd, Anomalous preasymptotic colloid transport by hydrodynamic dispersion in microfluidic capillary flow, Phys. Rev. E 90 (2014) 010301 (Jul 2014).

[61] K. N. Magdoom, A. Zeinomar, R. R. Lonser, M. Sarntinoranont, T. H. Mareci, Phase contrast mri of creeping flows using stimulated echo, Journal of Magnetic Resonance 299 (2019) 49–58 (2019).

[62] A. Vallatos, H. F. Al-Mubarak, J. M. Mullin, W. M. Holmes, Accuracy of phase-contrast velocimetry in systems with skewed intravoxel velocity distributions, Journal of Magnetic Resonance 296 (2018) 121–129 (2018).

[63] T. E. Conturo, G. D. Smith, Signal-to-noise in phase angle reconstruction: dynamic range extension using phase reference offsets, Magnetic Resonance in Medicine 15 (3) (1990) 420–437 (1990).

[64] D. Giese, M. Haeberlin, C. Barmet, K. P. Pruessmann, T. Schaeffter, S. Kozerke, Analysis and correction of background velocity offsets in phase-contrast flow measurements using magnetic field monitoring, Magnetic resonance in medicine 67 (5) (2012) 1294–1302 (2012).

[65] J. Busch, S. J. Vannesjo, C. Barmet, K. P. Pruessmann, S. Kozerke, Analysis of temperature dependence of background phase errors in phase-contrast cardiovascular magnetic resonance, Journal of Cardiovascular Magnetic Resonance 16 (1) (2014) 97 (2014).

[66] M. A. Bernstein, X. J. Zhou, J. A. Polzin, K. F. King, A. Ganin, N. J. Pelc, G. H. Glover, Concomitant gradient terms in phase contrast mr: Analysis and correction, Magnetic Resonance in Medicine 39 (2) (1998) 300–308 (1998).

[67] P. G. Walker, G. B. Cranney, M. B. Scheidegger, G. Waseleski, G. M. Pohost, A. P. Yoganathan, Semiautomated method for noise reduction and background phase error correction in mr phase velocity data, Journal of Magnetic Resonance Imaging 3 (3) (1993) 521–530 (1993).

[68] M. E. Komlosh, D. Benjamini, N. Williamson, F. Horkay, E. B. Hutchinson, P. J. Basser, A novel mri phantom to study interstitial fluid transport in the glymphatic system, Magnetic resonance imaging 56 (2019) 181–186 (2019).

[69] L. Lebon, L. Oger, J. Leblond, J. Hulin, N. Martys, L. Schwartz, Pulsed gradient nmr measurements and numerical simulation of flow velocity distribution in sphere packings, Physics of Fluids 8 (2) (1996) 293–301 (1996).

[70] S. L. Codd, B. Manz, J. D. Seymour, P. T. Callaghan, Taylor dispersion and molecular displacements in poiseuille flow, Phys. Rev. E 60 (1999) R3491–R3494 (Oct 1999).

[71] S. Stapf, K. Packer, R. Graham, J.-F. Thovert, P. Adler, Spatial correlations and dispersion for fluid transport through packed glass beads studied by pulsed field-gradient nmr, Physical Review E 58 (5) (1998) 6206 (1998).

[72] U. M. Scheven, P. N. Sen, Spatial and temporal coarse graining for dispersion in randomly packed spheres, Physical review letters 89 (25) (2002) 254501 (2002).

[73] A. Khrapitchev, P. Callaghan, Reversible and irreversible dispersion in a porous medium, Physics of Fluids 15 (9) (2003) 2649–2660 (2003).

[74] U. Scheven, R. Harris, M. Johns, Intrinsic dispersivity of randomly packed monodisperse spheres, Physical review letters 99 (5) (2007) 054502 (2007).

[75] M. Hunter, P. Callaghan, Nmr measurement of nonlocal dispersion in complex flows, Physical review letters 99 (21) (2007) 210602 (2007).

[76] D. Le Bihan, E. Breton, D. Lallemand, P. Grenier, E. Cabanis, M. Laval-Jeantet, Mr imaging of intravoxel incoherent motions: application to diffusion and perfusion in neurologic disorders., Radiology 161 (2) (1986) 401–407 (1986).

[77] D. Le Bihan, E. Breton, D. Lallemand, M. Aubin, J. Vignaud, M. Laval-Jeantet, Separation of diffusion and perfusion in intravoxel incoherent motion mr imaging., Radiology 168 (2) (1988) 497–505 (1988).

[78] T. Taoka, Y. Masutani, H. Kawai, T. Nakane, K. Matsuoka, F. Yasuno, T. Kishimoto, S. Naganawa, Evaluation of glymphatic system activity with the diffusion mr technique: diffusion tensor image analysis along the perivascular space (dti-alps) in alzheimer’s disease cases, Japanese journal of radiology 35 (4) (2017) 172–178 (2017).

[79] J. J. Iliff, M. Wang, D. M. Zeppenfeld, A. Venkataraman, B. A. Plog, Y. Liao, R. Deane, M. Nedergaard, Cerebral arterial pulsation drives paravascular csf–interstitial fluid exchange in the murine brain, Journal of Neuroscience 33 (46) (2013) 18190–18199 (2013).

[80] C. W. Windt, F. J. Vergeldt, P. A. De Jager, H. Van As, Mri of long-distance water transport: a comparison of the phloem and xylem flow characteristics and dynamics in poplar, castor bean, tomato and tobacco, Plant, Cell & Environment 29 (9) (2006) 1715–1729 (2006).

[81] A. Colbourne, A. Sederman, M. Mantle, L. Gladden, Accelerating flow propagator measurements for the investigation of reactive transport in porous media, Journal of Magnetic Resonance 272 (2016) 68–72 (2016).

[82] D. Benjamini, M. E. Komlosh, N. H. Williamson, P. J. Basser, Generalized mean apparent propagator mri to measure and image advective and dispersive flows in medicine and biology, IEEE transactions on medical imaging 38 (1) (2018) 11–20 (2018).

[83] D. Bourgeois, M. Decorps, Quantitative imaging of slow coherent motion by stimulated echoes with suppression of stationary water signal, Journal of Magnetic Resonance (1969) 94 (1) (1991) 20–33 (1991).

[84] T. Scheenen, F. Vergeldt, C. Windt, P. De Jager, H. Van As, Microscopic imaging of slow flow and diffusion: a pulsed field gradient stimulated echo sequence combined with turbo spin echo imaging, Journal of Magnetic Resonance 151 (1) (2001) 94–100 (2001).

[85] A. Ahlgren, L. Knutsson, R. Wirestam, M. Nilsson, F. Ståhlberg, D. Topgaard, S. Lasič, Quantification of microcirculatory parameters by joint analysis of flow-compensated and non-flow-compensated intravoxel in-coherent motion (ivim) data, NMR in Biomedicine 29 (5) (2016) 640–649 (2016).

[86] R. Mills, Self-diffusion in normal and heavy water in the range 1-45. deg., The Journal of Physical Chemistry 77 (5) (1973) 685–688 (1973).

[87] C. L. Dumoulin, S. P. Souza, R. D. Darrow, N. J. Pelc, W. J. Adams, S. A. Ash, Simultaneous acquisition of phase-contrast angiograms and stationary-tissue images with hadamard encoding of flow-induced phase shifts, Journal of Magnetic Resonance Imaging 1 (4) (1991) 399–404 (1991).

[88] H. Lee, K. Mortensen, S. Sanggaard, P. Koch, H. Brunner, B. Quistorff, M. Nedergaard, H. Benveniste, Quantitative gd-dota uptake from cerebrospinal fluid into rat brain using 3d vfa-spgr at 9.4 t, Magnetic resonance in medicine 79 (3) (2018) 1568–1578 (2018).

[89] L. A. Ray, J. J. Heys, Fluid flow and mass transport in brain tissue, Fluids 4 (4) (2019) 196 (2019).

[90] L. Xie, H. Kang, Q. Xu, M. J. Chen, Y. Liao, M. Thiyagarajan, J. O’Donnell, D. J. Christensen, C. Nicholson, J. J. Iliff, et al., Sleep drives metabolite clearance from the adult brain, science 342 (6156) (2013) 373–377 (2013).

[91] S. Kounda, R. Elkin, S. Nadeem, Y. Xue, S. Constantinou, S. Sanggaard, X. Liu, B. Monte, F. Xu, W. Van Nostrand, et al., Optimal mass transport with lagrangian workflow reveals advective and diffusion driven solute transport in the glymphatic system, Scientific reports 10 (1) (2020) 1–18 (2020).

[92] L. Ray, J. J. Iliff, J. J. Heys, Analysis of convective and diffusive transport in the brain interstitium, Fluids and Barriers of the CNS 16 (1) (2019) 6 (2019).

[93] N. Nishimura, C. B. Schaffer, B. Friedman, P. D. Lyden, D. Kleinfeld, Penetrating arterioles are a bottleneck in the perfusion of neocortex, Proceedings of the National Academy of Sciences 104 (1) (2007) 365–370 (2007).

[94] D. L. Adams, V. Piserchia, J. R. Economides, J. C. Horton, Vascular supply of the cerebral cortex is specialized for cell layers but not columns, Cerebral Cortex 25 (10) (2014) 3673–3681 (2014).

[95] M. L. Rennels, T. F. Gregory, O. R. Blaumanis, K. Fujimoto, P. A. Grady, Evidence for a ‘paravascular’fluid circulation in the mammalian central nervous system, provided by the rapid distribution of tracer protein throughout the brain from the subarachnoid space, Brain research 326 (1) (1985) 47–63 (1985).

[96] K. N. Magdoom, A. Brown, J. Rey, T. H. Mareci, M. A. King, M. Sarntinoranont, Mri of whole rat brain perivascular network reveals role for ventricles in brain waste clearance, Scientific reports 9 (1) (2019) 1–11 (2019).

[97] H. F. Cserr, L. Ostrach, Bulk flow of interstitial fluid after intracranial injection of blue dextran 2000, Experimental neurology 45 (1) (1974) 50–60 (1974).

[98] H. Mestre, J. Tithof, T. Du, W. Song, W. Peng, A. M. Sweeney, G. Olveda, J. H. Thomas, M. Nedergaard, D. H. Kelley, Flow of cerebrospinal fluid is driven by arterial pulsations and is reduced in hypertension, Nature communications 9 (1) (2018) 4878 (2018).

[99] B. Bedussi, M. Almasian, J. de Vos, E. VanBavel, E. N. Bakker, Paravascular spaces at the brain surface: Low resistance pathways for cerebrospinal fluid flow, Journal of Cerebral Blood Flow & Metabolism 38 (4) (2018) 719–726 (2018).

[100] F. Sepehrband, G. Barisano, N. Sheikh-Bahaei, R. P. Cabeen, J. Choupan, M. Law, A. W. Toga, Image processing approaches to enhance perivascular space visibility and quantification using mri, Scientific reports 9 (1) (2019) 1–12 (2019).

