## Supplementary figures S1--S11 for "Limits to flow detection in phase contrast MRI"

Supplementary figures S1–S5 show axial beadpack, axial bulk, or sagittal images of phase and magnitude at $\pm q_4$ and the difference between the $+q_4$ and $-q_4$ for flow rates $\dot{\mathrm{V}} = 100$ or $0\ \mu$l/min. The captions of Figs. S3–S5 are reduced for brevity but are similar to the Fig. S2 caption. The diagonal stripes in the axial phase images were found to be affected by EPI double sampling and are therefore likely a consequence of the EPI readout. A hot spot is seen in the upper right corner of axial beadpack phase difference images and magnitude images. Since it is also seen at $\dot{\mathrm{V}} = 0$, it is likely an artifact. These figures supplement Fig. 4 in the paper.

Fig. S6 shows axial images of the phase difference at $\dot{\mathrm{V}} = 0$ for all $q$ values. Note that the scale bar is enhanced to show detail. Also note that the phase difference should be zero if $v = 0$ and any non-zero phase shift leads to a measured velocity offset. Fig. S7 was made in the same way, but for $\dot{\mathrm{V}} = 300\ \mu$l/min. This is well into the fast-flow regime. As a consequence of $q$ being larger than optimal, flow aliasing is seen in the regions with highest velocity for $q_3$, $q_4$, and $q_5$.

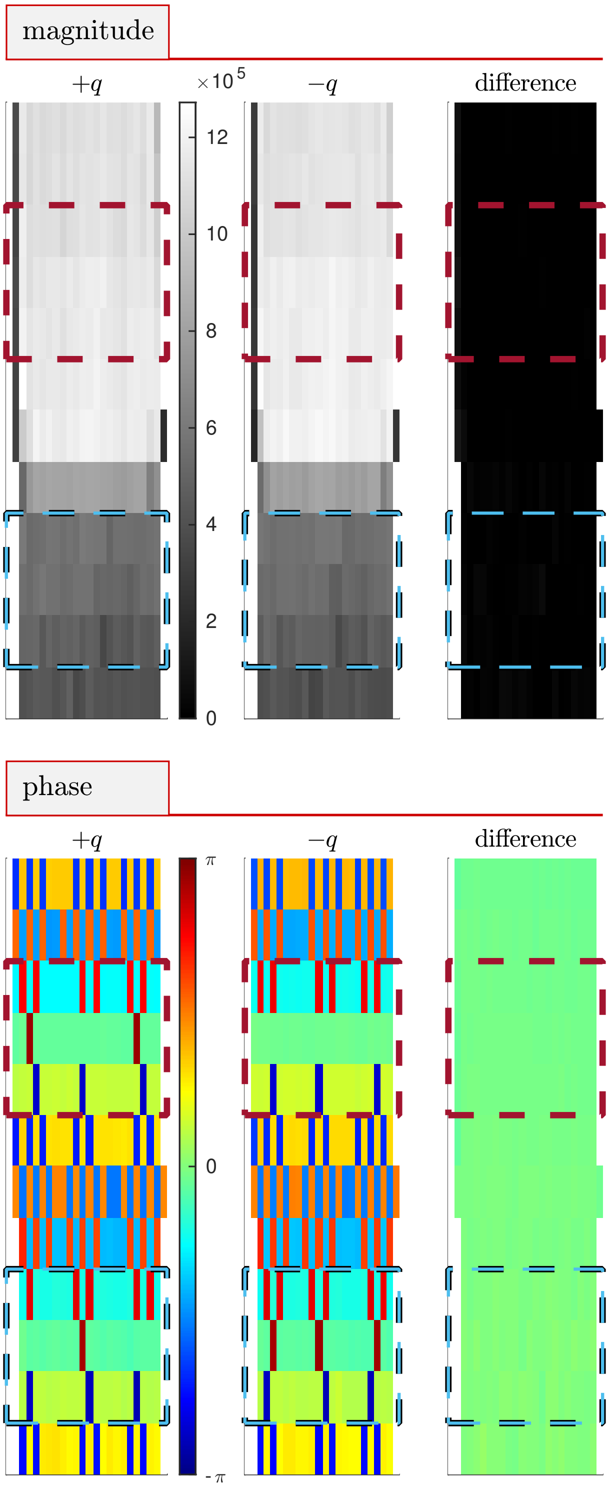


Figure S1: **Magnitude and phase images, $\dot{\text{V}} = 0$.** Sagittal images of magnitude and phase at $+q_4$, $-q_4$ and the difference between $+q_4$ and $-q_4$. Figure accompanies Fig. 4 in the paper.

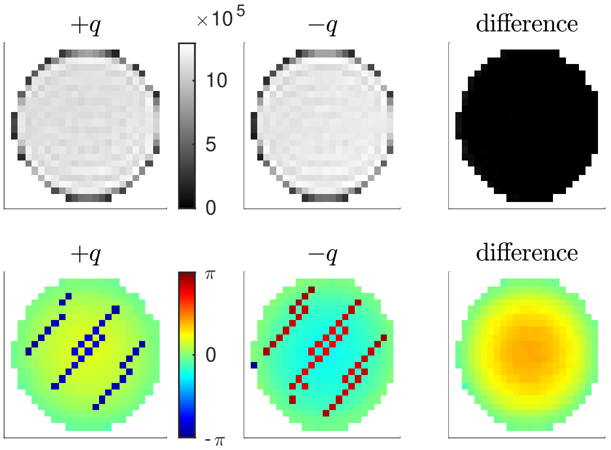


Figure S2: **Magnitude and phase images, bulk slice, with flow.** Axial images of magnitude (top) and phase (bottom) at $+q_4$, $-q_4$ and the difference between $+q_4$ and $-q_4$ for the bulk region with $\dot{\text{V}} = 100$ $\mu$l/min.

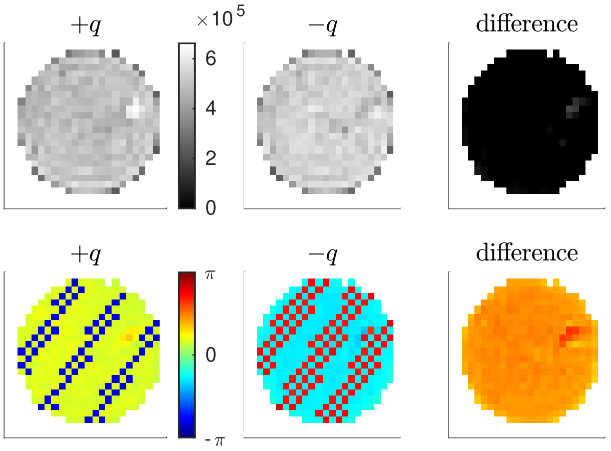


Figure S3: **Magnitude and phase images, beadpack slice, $\dot{\text{V}} = 100$ $\mu$l/min, $q_4$.**

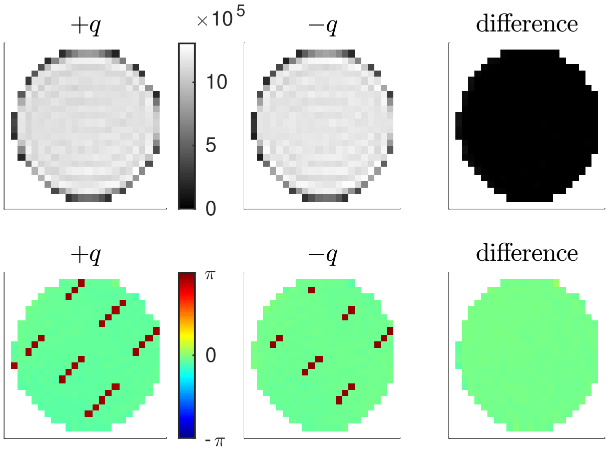


Figure S4: **Magnitude and phase images, bulk slice, $\dot{\text{V}} = 0$, $q_4$.**

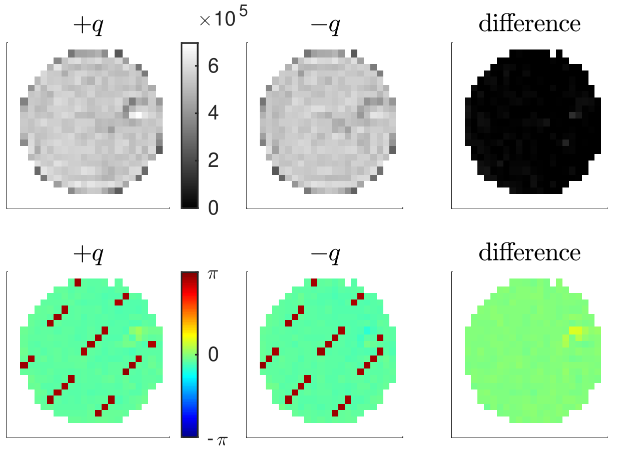


Figure S5: **Magnitude and phase images, beadpack slice, $\dot{V}$ = 0, $q_4$.**

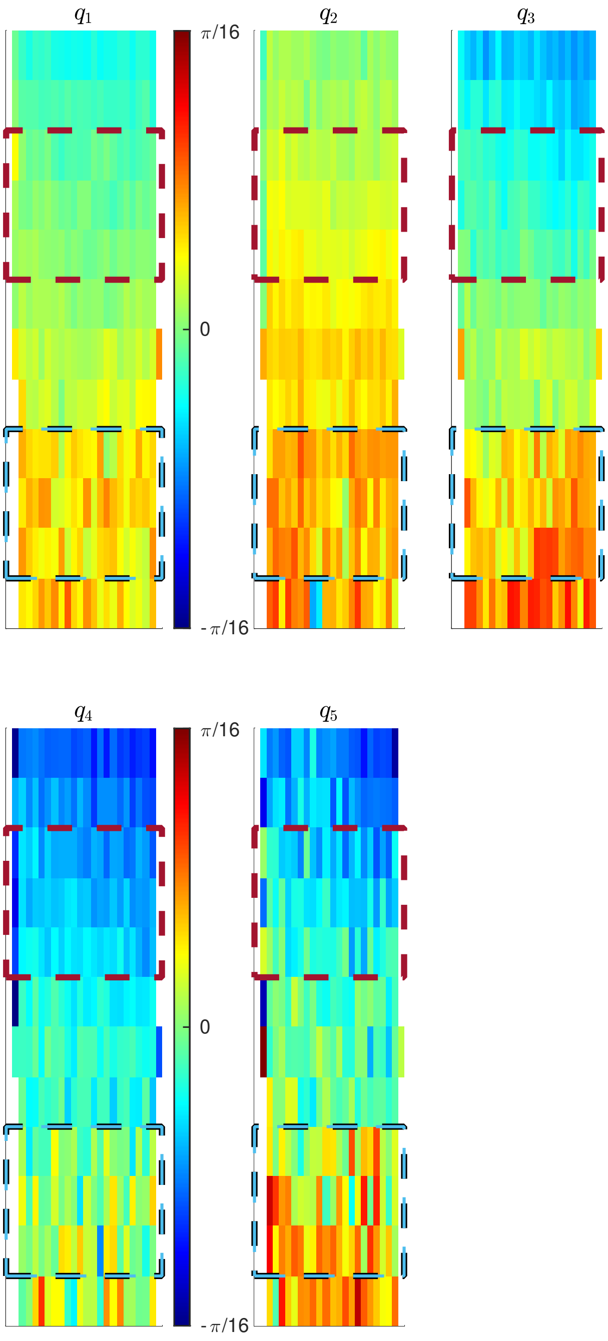


Figure S6: **Phase difference images.** Sagittal images of the phase difference between $+q$ and $-q$ for $\dot{V}$ = 0 at different $q$-points. Note the different scale of the color bar.

Fig. S8 shows velocity images at the same flow rates as presented in Fig. 5, but after subtracting the phase phase offset measured at the $\dot{V} = 0$. Fig. 5 can be referenced for the full figure caption.

Fig. S9 shows velocity images at select flow rates in the fast-flow regime using $q_2$. No phase wrapping (flow aliasing) is observed at this q-value. Slower velocity at the outer radius of the beadpack is observed at these faster flows, but is not evident at slower flows. Additionally, the velocity profile in axial slices through the bulk region just above the beadpack no longer appear parabolic. A blunted profile expected at these higher velocities, since hydrodynamic entrance length effects scale with Reynolds number (which is proportional to $v$).

Fig. S10 compares the experimentally measured velocities and actual velocities for all $q$ values and supplements Fig. 6 in the paper. The velocity offset at $\dot{V} = 0$ is apparent, particularly at $q_1$–$q_3$. The transition from unmeasurable to measurable velocity is similar for all $q$-values. In the fast velocity region, flow aliasing wraps the $q_4$ and $q_5$ velocity measurements to lower values (as well as $q_3$ for the fastest flow rate). Most importantly, the measured velocities are consistent with the actual velocities.

Fig. S11 compares the experimentally measured flow rates and actual flow rates for all $q$ values and supplements Fig. 7. The experimentally measured flow rate is the average velocity in a slice divided by the area which is open to flow. For the bulk region this is the cross-sectional area. For the beadpack it is the cross-sectional area multiplied by the porosity. The actual flow rate is defined by the pump. By mass conservation, the flow rate through all slices should be equal to the actual flow rate. For the beadpack, the data differs from fig. S10 because here the standard deviation is taken over the three slices whereas in fig. S10 the standard deviation is taken over all velocities. Again, the offset at $\dot{V} = 0$ is apparent and varies with $q$ and location within the sample.

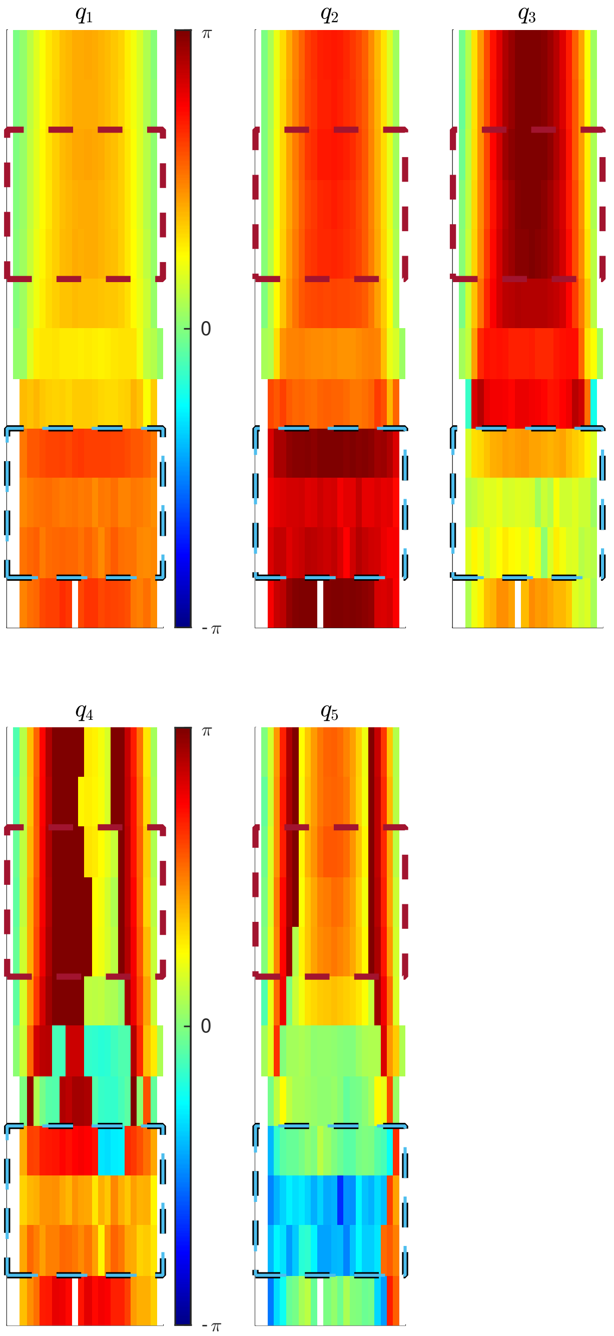


Figure S7: **Phase difference images, fast flow.** Sagittal images of the phase difference between $+q$ and $-q$ for $\dot{V} = 300\,\mu$l/min. Note wrapping backwards by $\pi$.

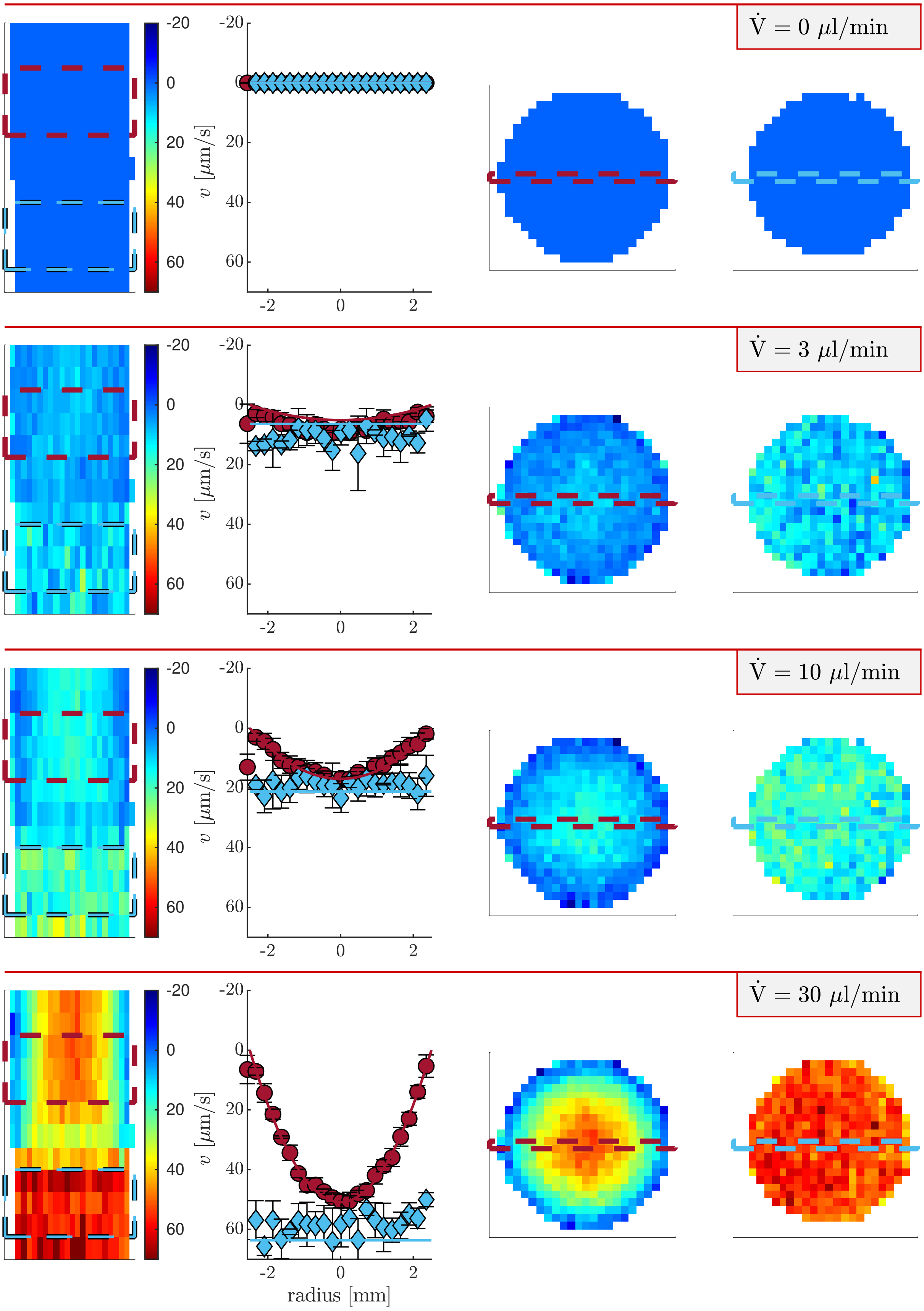


Figure S8: **Velocity images and profiles of the beadpack flow phantom at slow flows using** $q_4$**.** The phase image acquired at $\dot{V}$ = 0 is subtracted to remove the velocity offset.

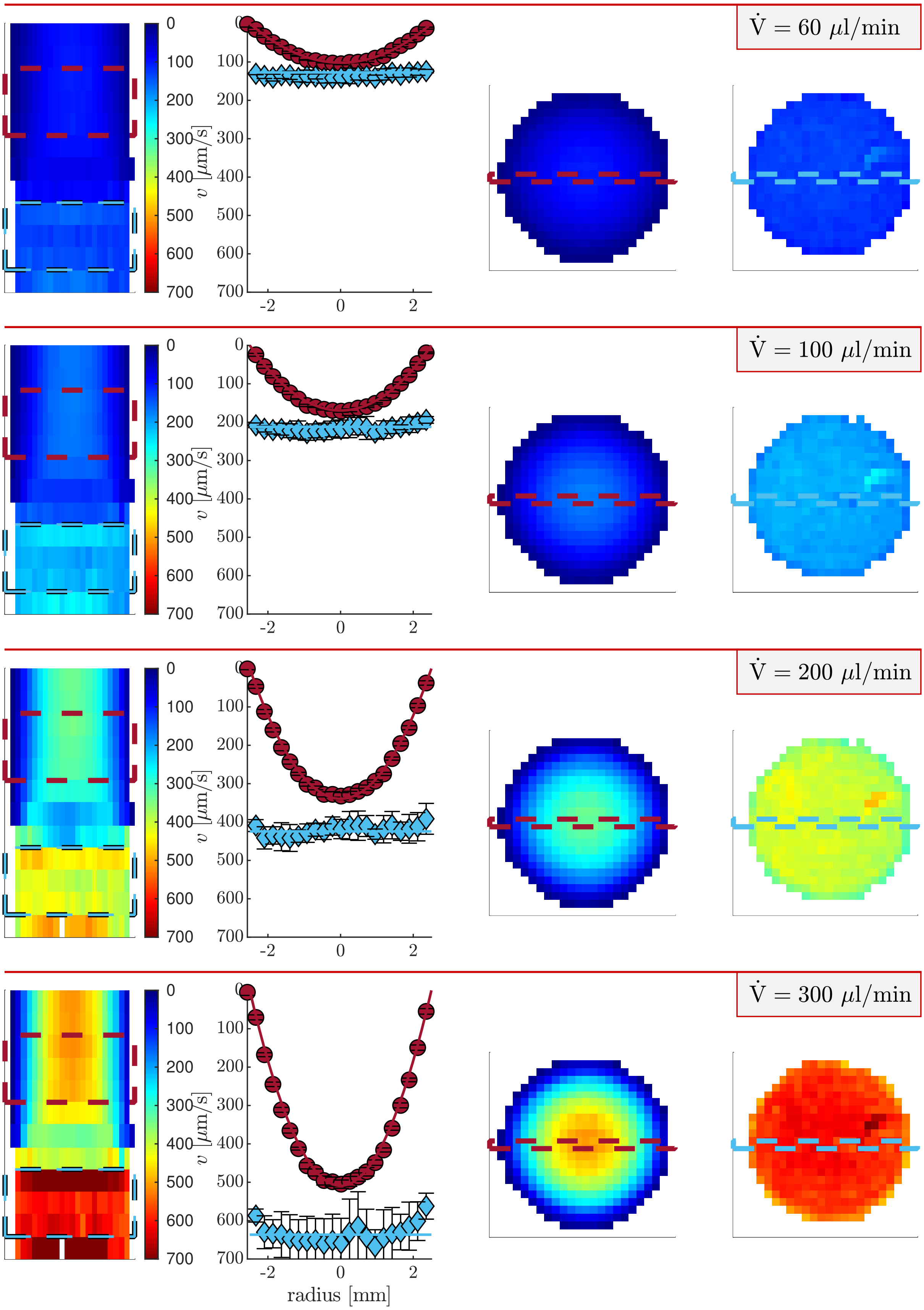


Figure S9: **Velocity images and profiles of the beadpack flow phantom at the transition from slow- to fast-flow regimes using $q_2$.** $\dot{V}$ = 300 crosses into the fast-flow regime. $q_2 < q_{slow}$ and as such is near-optimal.

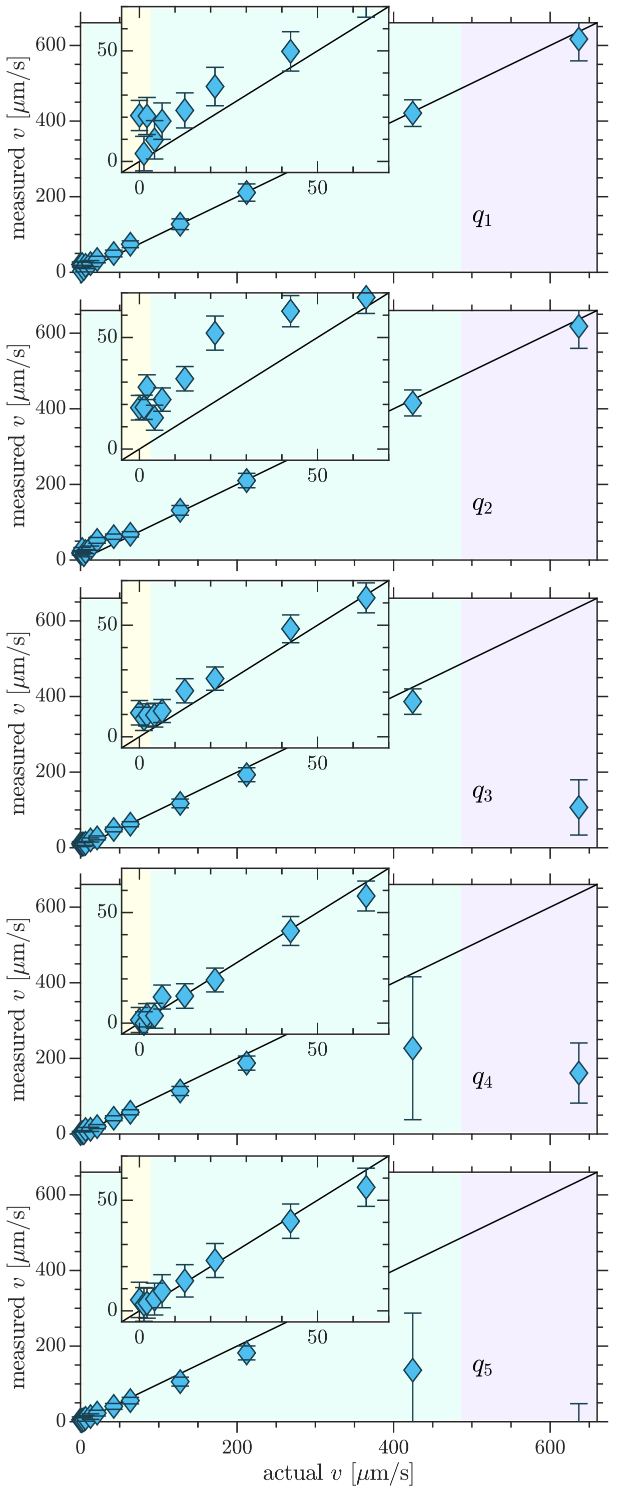


Figure S10: **Experimental limits to flow detection with phase contrast velocimetry.** Comparison of experimentally measured and actual velocities for flow through the beadpack at different $q$-points. See Fig. 6 for full figure caption.

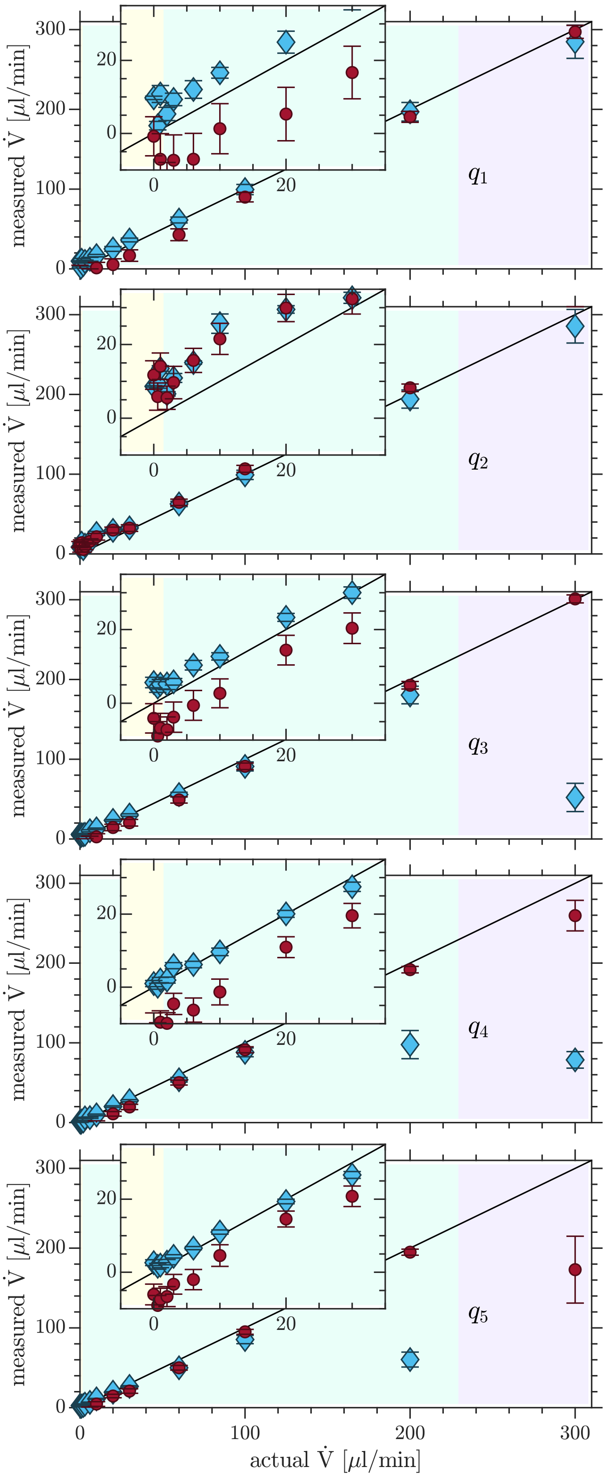


Figure S11: **Measured vs. actual flow rate.** Means and standard deviations are from the flow rates measured through three slices for both the beadpack (blue diamonds) and bulk (red circles) ROIs, shown in Fig. 4 and other images.
